# Pangenome-based genome inference

**DOI:** 10.1101/2020.11.11.378133

**Authors:** Jana Ebler, Wayne E. Clarke, Tobias Rausch, Peter A. Audano, Torsten Houwaart, Jan Korbel, Evan E. Eichler, Michael C. Zody, Alexander T. Dilthey, Tobias Marschall

## Abstract

Typical analysis workflows map reads to a reference genome in order to detect genetic variants. Generating such alignments introduces references biases, in particular against insertion alleles absent in the reference and comes with substantial computational burden. In contrast, recent k-mer-based genotyping methods are fast, but struggle in repetitive or duplicated regions of the genome. We propose a novel algorithm, called PanGenie, that leverages a pangenome reference built from haplotype-resolved genome assemblies in conjunction with k-mer count information from raw, short-read sequencing data to genotype a wide spectrum of genetic variation. The given haplotypes enable our method to take advantage of linkage information to aid genotyping in regions poorly covered by unique k-mers and provides access to regions otherwise inaccessible by short reads. Compared to classic mapping-based approaches, our approach is more than 4× faster at 30× coverage and at the same time, reached significantly better genotype concordances for almost all variant types and coverages tested. Improvements are especially pronounced for large insertions (> 50bp), where we are able to genotype > 99.9% of all tested variants with over 90% accuracy at 30× short-read coverage, where the best competing tools either typed less than 60% of variants or reached accuracies below 70%. PanGenie now enables the inclusion of this commonly neglected variant type in downstream analyses.

## 1 Introduction

Diploid organisms have two copies of each autosomal chromosome, each of which can carry genetic variation. The process of determining whether a known variant allele is located on one or both of these copies, or whether the variant is absent in an individual’s genome, is referred to as *genotyping*. Different classes of genetic variants exist and include SNPs (single nucleotide polymorphisms), indels (insertions and deletions) up to 50 bp in size, and larger structural variants (SVs). Large studies have produced comprehensive catalogues of known variation of various types, ranging from single-nucleotide variants (SNVs) to large structural variants (SVs), for the human genome [1, 2, 3, 4]. Many variants have been linked to diseases, such as schizophrenia or autism, which makes genotyping an essential task for studying such diseases [5, 6, 7, 8, 9, 10]. Widely used genotyping methodss, such as GATK [11], FreeBayes [12], Delly [13], Platypus [14] and SVTyper [15], are based on short-read alignments to a reference genome and thus, come with a reference bias, as the aligner is unaware of possible alternative sequences that might be present in an individual’s genome [16, 17]. This can be especially problematic when genotyping structural variants, defined as events of 50bp and longer. Recently, several approaches have been suggested that replace the linear reference genome by graph structures which include possible alternative alleles. Graphs are either built from given variant calls or haplotype-resolved assemblies, and genotypes are derived from alignments of sequencing reads to these graphs [18, 17, 19, 20, 21]. In general, these graph-based approaches were shown to improve genotyping accuracy over methods that rely on a linear reference genome by reducing reference bias. However, aligning sequencing reads is a time consuming task even for linear reference genomes, where mapping 30× short read sequencing data of a single sample takes around 100 CPU hours. This problem is amplified when transitioning to graph-based pangenome references, where the read mapping problem is even more computationally expensive.

A much faster alternative is to genotype known variants based on counts of *k*-mers, short sequences of a fixed length *k*, in the raw sequencing reads. Cortex [22] was the first approach to genotyping variants leveraging read *k*-mer count information based on a colored de Bruijn graph constructed from sequencing data and known allelic sequences. Dilthey et al. [23] use a similar idea and construct population reference graphs from known haplotype sequences to genotype a sample’s MHC region based on short read sequencing data, but this approach does not scale to whole genomes.

Dolle et al. [24] genotype SNPs and short indels based on querying reads containing allele specific k-mers in the data. They derive genotypes from alignments of these reads to reference and alternative sequences. BayesTyper [25] constructs graphs containing reference and alternative alleles for sets of variants that are less than a *k*-mer size apart in the genome and genotypes are computed by sampling the likeliest pair of local haplotypes through each such cluster of variants, based on the observed k-mer count profiles. Such *k*-mer-based methods allow fast genotyping by bypassing the time consuming alignment step. However, they can struggle in repetitive and duplicated regions of the genome which are not covered by any unique *k*-mers, as they lack the connectivity information contained in the reads. This is especially problematic for structural variants which are often located in repeat-rich or duplicated regions of the genome [26, 3].

Turner et al. [27] aim to address this problem by introducing linked de Bruijn graphs which store long range connectivity information from sequencing reads on top of a de Bruijn graph. They demonstrated that adding link information from a set of reference sequences to the graph in this way improved drug resistance locus assembly in *K*.*pneumoniae* isolates. In a similar manner, information of already known haplotype sequences of other samples could improve *k*-mer-based genotyping especially in difficult to access regions of large diploid genomes, but methods for this have so far been lacking. Known haplotypes (in form of a reference panel) have been used previously for population based phasing of small variants. The Li-Stephens Model provides a theoretical framework by formulating this problem in terms of a Hidden Markov Model [28]. Furthermore, reference panel information can be used to impute missing genotypes of a sample [29, 30, 31, 32], but accurate SV-integrated reference panels have been challenging to construct.

Recently, single molecule sequencing technologies delivering long read data have enabled breakthroughs in producing *de novo* haplotype-resolved genome assemblies [33, 34, 35]. Such assemblies are already available for several human samples and major efforts are underway^1^ to generate hundreds of human genome assemblies with the intention of deriving a pangenome representation that replaces the current reference genome GRCh38. So far, however, scalable methods to leverage such haplotype-resolved pangenome representations for the interpretation of short-read data sets are not available.

In this paper, we describe an algorithm, PanGenie (for *Pangenome-based Genome Inference*), that makes use of haplotype information from an assembly-derived pangenome representation in combination with read *k*-mer counts for efficiently genotyping a wide spectrum of variants. That is, our method is able to leverage the information inherent in the assemblies in order to infer the genome of a new sample for which only short-reads are available. PanGenie bypasses read-mapping and is entirely based on *k*-mers, which allows it to rapidly proceed from the input short reads to a final call set including SNPs, indels and structural variants, enabling access to variants typically not accessible in short-read workflows – such as larger insertions. We applied our method to genotype variants called from haplotype-resolved assemblies of six individuals, revealing a substantial advance in terms of runtime, genotyping accuracy, and in the number of accessible variants.

## 2 Results

### 2.1 Algorithm overview

The input to our algorithm consists of short read sequencing data for the sample to be genotyped, a reference genome, as well as a pangenome graph containing variants and paths representing known haplotype sequences. The latter is represented in terms of a fully-phased, multisample VCF file. In a first step, clusters of variants less than the k-mer size apart are combined into single, multi-allelic variants. We identify all k-mers unique to a variant region, that is, k-mers that cover the variant position and do not occur anywhere else in the genome, and use Jellyfish [36] to determine their counts in the reads. Our genotyping model combines two sources of information in order to derive genotypes for the variants: read k-mer counts and the already known haplotype sequences. The distribution of k-mer counts along the allele paths of a variant can hint towards the genotype of the sample. Figure 1a provides an example: three alleles are shown for a variant. All k-mers corresponding to the middle one are absent from the reads of the sample. This indicates that the individual carries the red and the blue allele at this position. However, variants may be poorly covered by k-mers, or no unique k-mers may exist for a variant in repetitive regions of the genome. Such positions cannot be reliably genotyped by an approach based purely on the k-mer counts. In these regions, information of known haplotype sequences of a population can help to infer genotypes based on neighboring variants. An example is provided in Figure 1b: known haplotype sequences can be represented as paths in the graph. The second variant is poorly covered by k-mers but the count distribution of k-mers along the alleles of the first variant indicates that the unknown genome is composed of the green and blue haplotype.

**Figure 1:**
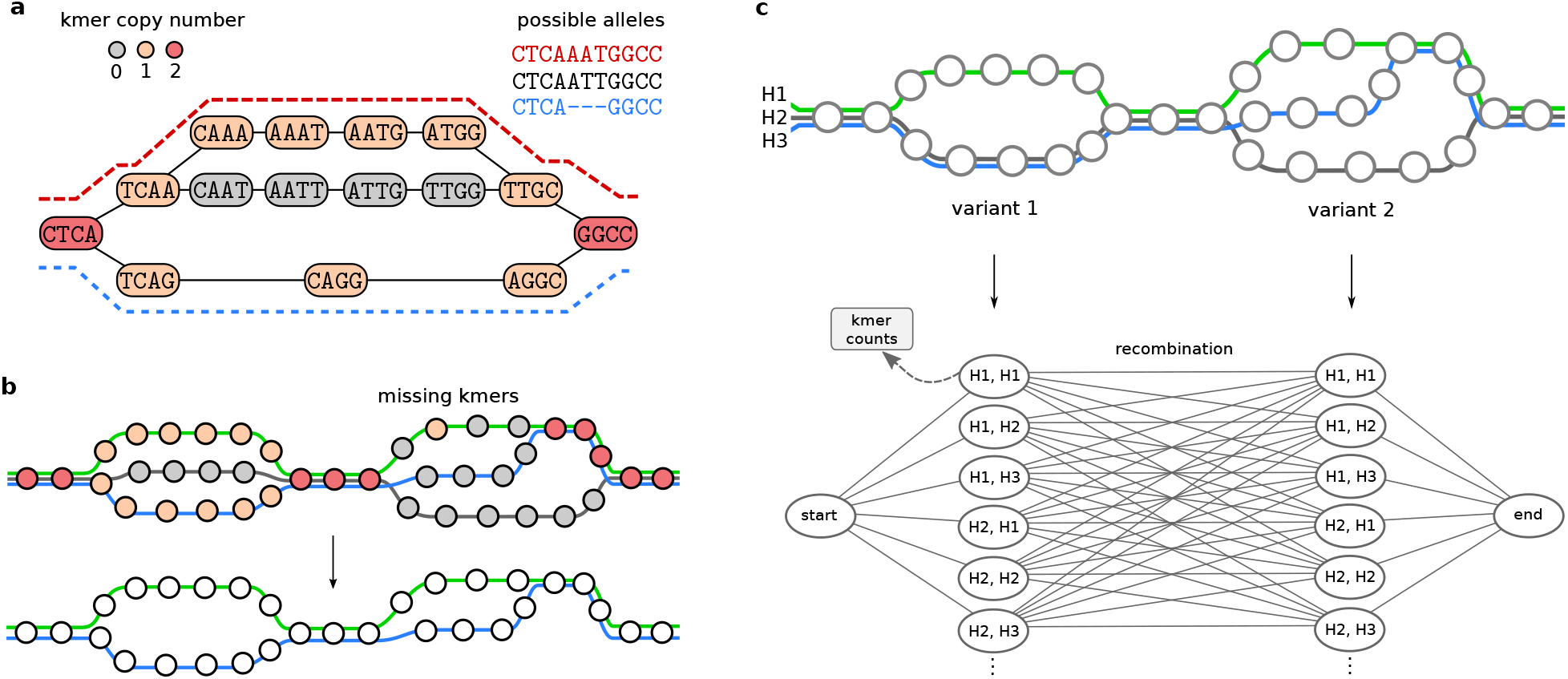
Genotyping approach. Our genotyping algorithm combines two sources of information: read k-mer counts and known haplotypes. **a)** A variant region in the constructed genome graph is shown. Each path corresponds to an allele. Colors indicate copy number estimates for the k-mers, based on which a genotype can be determined. Here, the variant likely carries the red and the blue allele, indicated by the two dashed lines. **b)** A larger proportion of the graph is shown, with three known haplotypes threaded through it. Again, colors indicate copy number estimates. The second bubble is poorly covered by k-mers, however, linkage to adjacent variants can be used to infer the two local haplotype paths. **c)** A genome graph with two variant positions is shown with the corresponding HMM below. Gray circles in the graph indicate k-mers. The hidden states of the HMM correspond to possible pairs of the three haplotype paths shown in the graph. These states output counts for unique k-mers characterizing the alleles.

For genotyping, we combine these two sources of information by constructing a Hidden Markov Model which models the unknown haplotypes of a sample as mosaics of the provided haplotypes and reconstructs them based on the read k-mer counts observed in the sample’s sequencing reads. To achieve this, our HMM has a hidden state for each possible pair of given haplotypes that can be chosen at each variant position. These states emit counts for the unique k-mers in the variant region based on the copy number of these k-mers in the two selected haplotypes. Changes in the selected haplotype paths between adjacent variant positions correspond to recombination events. Therefore, we define transition probabilities based on recombination probabilities defined in [28]. We show an example in Figure 1c. Running the Forward-Backward algorithm, we can compute genotype likelihoods for each position, from which we finally derive a genotype. Using the Viterbi algorithm, we can compute the two likeliest haplotype sequences given the observed k-mer counts.

### 2.2 Constructing a pangenome reference from haplotype-resolved assemblies

In order to construct a pangenome graph, we used haplotype-resolved assemblies of five individuals that have recently been produced [34, 35]. These samples include two individuals of Puerto Rican descent (HG00731, HG00732) as well as NA12878, NA24385 and PGP1. For each sample, we separately mapped contigs of each haplotype to the reference genome and used these alignments to call variants on each haplotype of all autosomes (see 4.1 for details). In order to filter out low quality or erroneous calls, we only kept variants located in regions in which all haplotypes were covered by exactly one contig alignment. These *callable* regions cover 87.42% (2.51 Gb) of chromosomes 1-22.

We create an acyclic and directed pangenome graph that contains bubbles representing the variation observed in all of the input haplotypes. Variants overlapping across haplotypes are combined into a single bubble with potentially multiple branches reflecting all allele sequences observed in the respective genomic region. The input haplotypes are represented as paths through the resulting pangenome. The final graph is represented in terms of a fully phased, multi-sample VCF file. Figure 2 provides an example of how we construct the graph.

**Figure 2:**
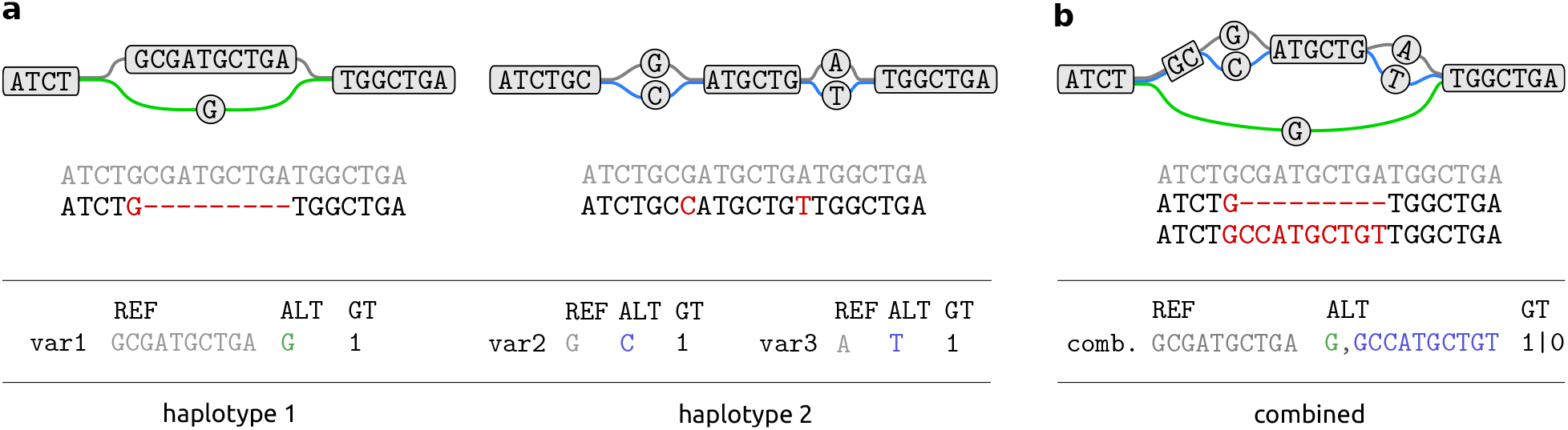
Combining variant calls. **a)** Shown are alignments of two contigs to the reference genome and variant calls for both of them: a deletion for the contig of the first haplotype, and two SNPs for the second haplotype. Additionally, the pangenome graphs that result from inserting these variants into the reference genome, are shown. **b)** Variant calls of both haplotypes are overlapping and will be represented as a single bubble when constructing a graph that contains all variants. In the resulting VCF, only those paths through the bubble will be listed, that were observed in at least one of the input haplotypes and additionally, the genotypes corresponding to each sample.

Due to the lack of haplotyped-resolved assemblies for other samples, the number of haplotype paths in our graph is relatively small. Until more assemblies become available in the future, we showcase the performance of our method by extending our pangenome reference panel using additional short read data sets. To this end, we apply PanGenie to phase the same set of variants in these additional samples. In this way, we used short-read sequencing data of a sample of Chinese descent, one individual of Yorubian descent as well as four samples from different populations (see Figure 3a and Section 4.1) in order to produce an “extended” panel consisting of eleven samples.

**Figure 3:**
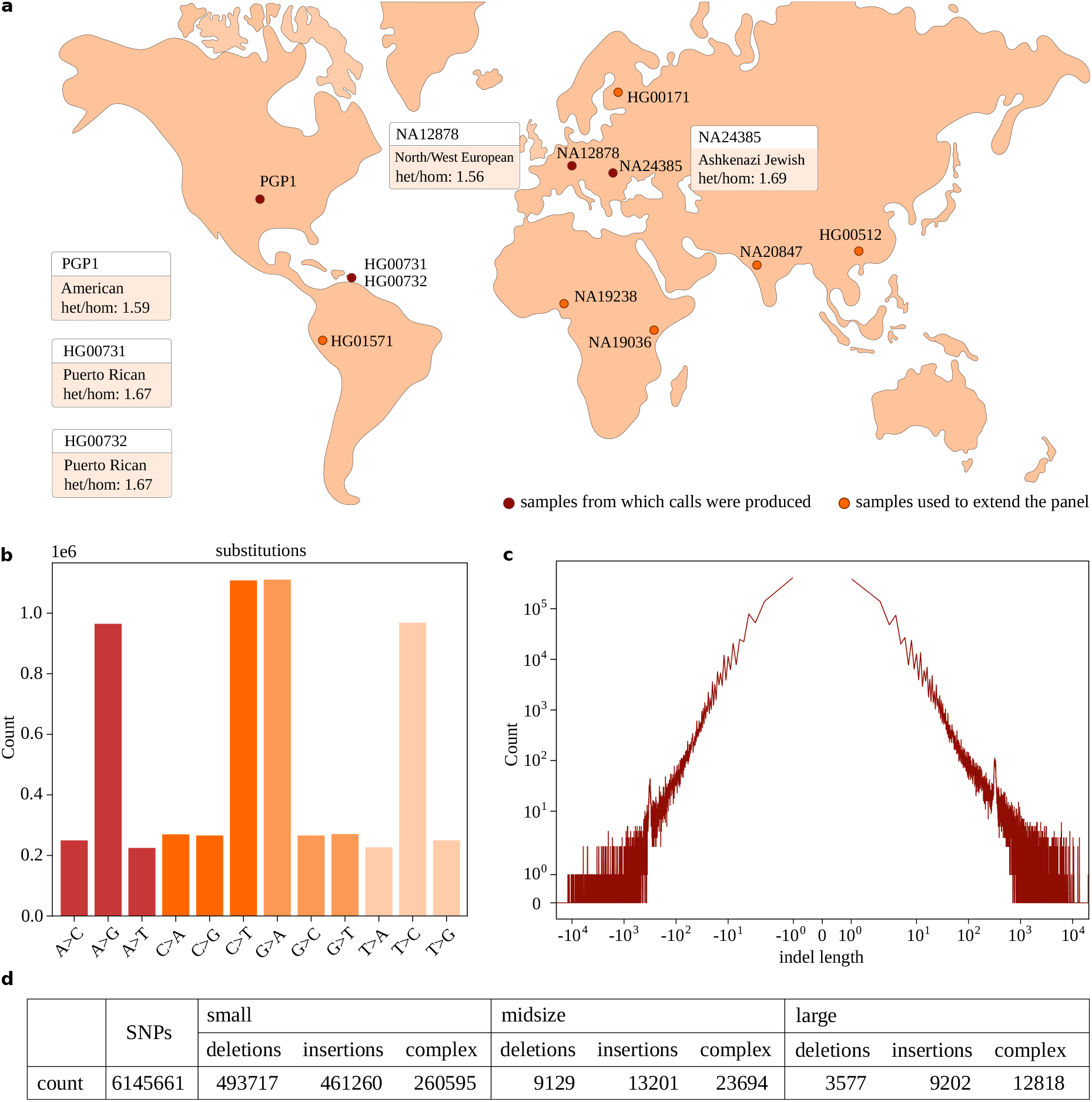
Callset statistics. Statistics for variants called from haplotype-resolved assemblies. **a)** Samples for which variants were called from haplotype-resolved assemblies are shown in red, as well as the population they originate from an the het/hom ratio observed for the variant calls. Furthermore, samples used to extend the panel are shown (orange). **b)** Shown are the number of different substitutions reported for all samples. **c)** Length distribution of insertions and deletions across all samples. Deletion lengths are reported as negative numbers, insertion lengths are positive. **d)** Number of variants per category: small (1 − 19bp), midsize (20 − 50bp), and large (> 50bp). Insertions and deletions include only bi-allelic variants, other types of structural variants or multi-allelic variants (that are no SNPs).

We present callset statistics in Figure 3. The transition/transversion (ti/tv) ratio for SNPs and the heterozygous/homozygous ratio are commonly used quality control measures for callsets [37, 38]. The ti/tv ratio is expected to be around two as transitions (changes from A to G, G to A, C to T and T to C) are twice as frequent as the remaining transversions. The distribution of these substitution types for our SNP calls are shown in in Figure 3b. We computed ti/tv ratios between 2.04 and 2.05 for all of our samples. Theoretically, the expected het/hom ratio for a callset is two for variants in Hardy-Weinberg equilibrium [37]. However, it has been previously reported to vary by ancestry and was observed to be smaller for individuals of American, Asian and European origin [38, 39]. This is in line with what we observe for our callsets. The five samples from which variant calls were generated are of American or European origin. We observed het/hom ratios between 1.56 and 1.67 for all these individuals (Figure 3a). Additionally, we show detailed counts observed for SNPs, insertions, deletions and complex variants of different lengths in Supplementary Table 2. Insertions and deletions include only bi-allelic variants, other types of structural variants or multi-allelic variants (that are not SNPs), are defined as complex variants. We distinguish small variants (1 − 19bp), midsize variants (20 − 50bp) and large variants (> 50bp). The total number of calls in each category can be found in Supplementary Table 3d. We additionally re-run our variant calling using reference version hg19 in order to be able to compare the resulting variant calls to the structural variants (> 50bp) contained in the Genome Aggregation Database (gnomAD) [4]. We found an overlap of 6,398 variants. 21,370 positions were only contained in our assembly-based callset set (see Supplementary Section 4.3).

### 2.3 Genotyping evaluation

For evaluation, we conducted a “leave-out-one experiment” and genotyped each of the four unrelated samples HG00731, HG00732, NA12878 and NA24385 based on Illumina reads from the HGSVC [3], the Genome in a Bottle Consortium [40] and 1000 Genomes Project high-coverage data (Mike Zody, personal communication). We used the variants from the pangenome graph constructed in Section 2.2 and two different sets of haplotype paths from this graph as input to our genotyping algorithm: first we only used the haplotypes that we obtained from haplotype-resolved assemblies, second, we added haplotypes from our extended callset. Thus, the former small panel consists of haplotypes for samples: HG00731, HG00732, NA12878, NA24385 and PGP1. The latter, extended panel, additionally contains haplotypes of: HG00512, NA19238, NA20847, NA19036, HG00171 and HG01571. We genotyped each of the four samples in a leave-one-out manner by removing it from the small and extended panels, respectively, and genotyped it based on the remaining samples. We then compare the genotype predictions to the ones of the left out, ground truth haplotypes derived from the haplotype-resolved assemblies. We additionally ran Platypus [14], BayesTyper [25], GATK HaplotypeCaller [11] and Paragraph [20] for comparison. Since Platypus, GATK and Paragraph are mapping-based approaches and require BAM-files as input, we used `bwa mem` [41] to align the reads to the reference genome prior to genotyping. PanGenie and BayesTyper are k-mer-based and were provided with the raw, unaligned sequencing reads (in FASTQ-format). We ran our experiments on different levels of read coverage. For this purpose we downsampled the reads of each sample to coverages 30×, 20×, 10× and 5×. Not all tools can handle all types of variants. We ran GATK only on SNPs, small and midsize variants and Paragraph was only run on midsize and large variants.

#### Evaluation metrics

Given a truth set of variants with known genotypes and genotype predictions made by a genotyper for these positions, we compute two metrics in order to evaluate the genotyping performance. The first one is the percentage of variants for which a tool was able to give a genotype prediction. Ideally, this fraction should be as large as possible. The second one is the *genotype concordance* that we define as the percentage of correct genotype predictions among all variants that were typed by the method.

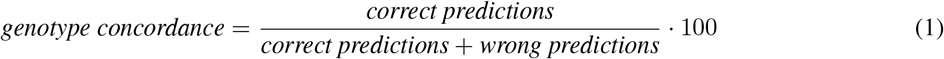

## Results

We show the genotyping results for sample HG00731 that we obtained from PanGenie using the extended panel as well as from the remaining methods in Figure 4 and 5. Respective results for the three other samples are similar and can be found in Supplementary Figures 8-13. The results we got from using the small panel are presented in Supplementary Figures 14-21. The plots show the genotype performances outside and inside of STR/VNTR regions, which we obtained from the UCSC genome browser [42]. We observed that between 54 − 78% of midsize and large variants are indeed located inside of repeats (Supplementary Table 3). Like most genotyping tools, PanGenie also calculates a phred-scaled genotype quality score which can be used to filter the genotypes. In our evaluation, we consider two configurations for PanGenie: “lenient” filtering, where we do not apply any filter and use all reported genotypes, and “strict”, where we only used high quality genotype predictions (quality >= 200) and treat all other variants as not genotyped. For all other tools, we did not apply any filters on the genotype quality and used all genotypes that they reported.

**Figure 4:**
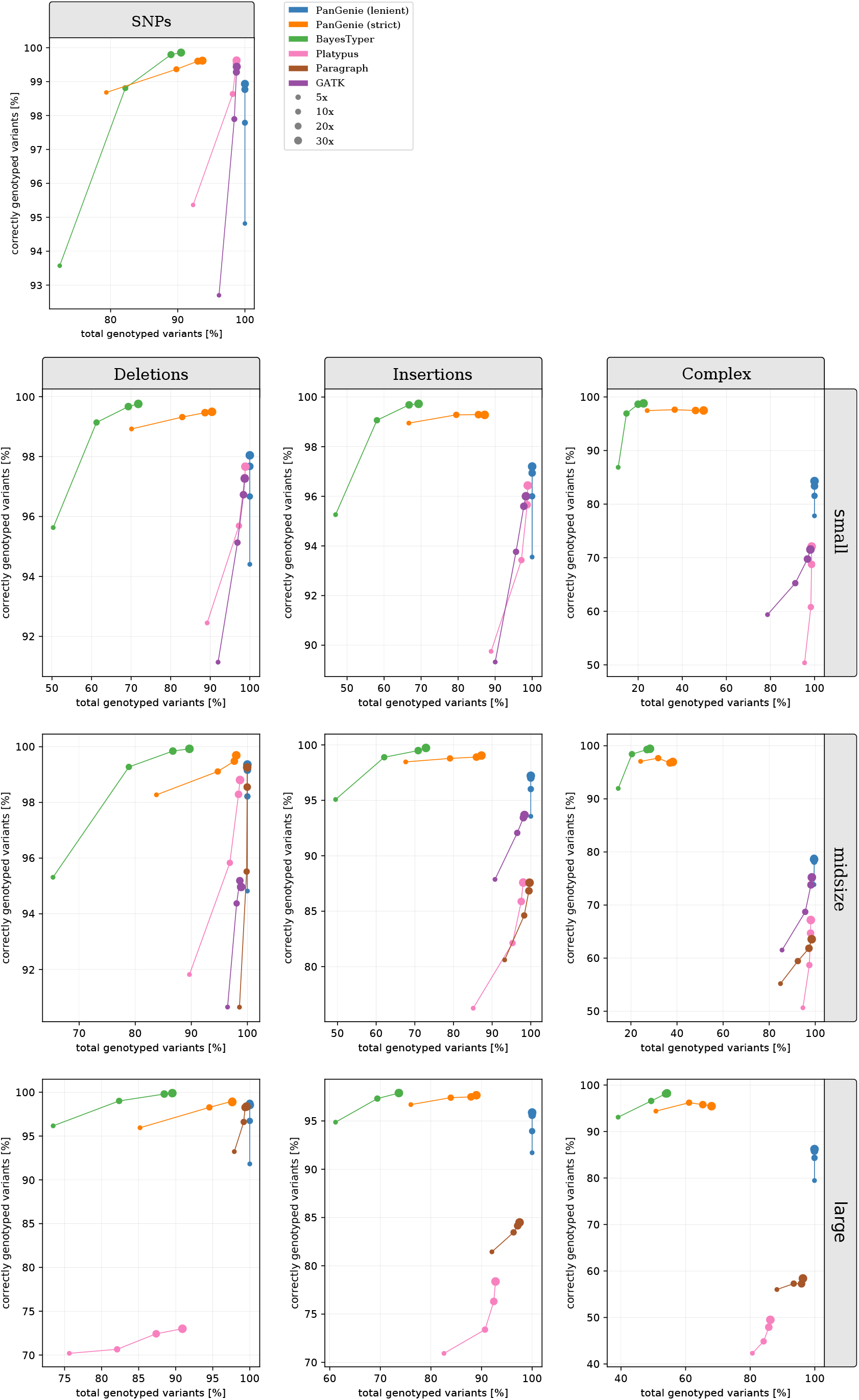
Non-repetitive regions. Genotype performance for sample **HG00731** at different coverages outside of STR/VNTR regions. We ran PanGenie using the **extended** panel, BayesTyper, Paragraph, Platypus and GATK in order to regenotype all callset variants. Besides not applying any filter on the reported genotype qualities (“lenient”), we additionally report genotyping statistics for PanGenie when using “strict” filtering (genotype quality >= 200). Note that GATK was not run on large variants, and Paragraph was only run on midsize and large variants.

**Figure 5:**
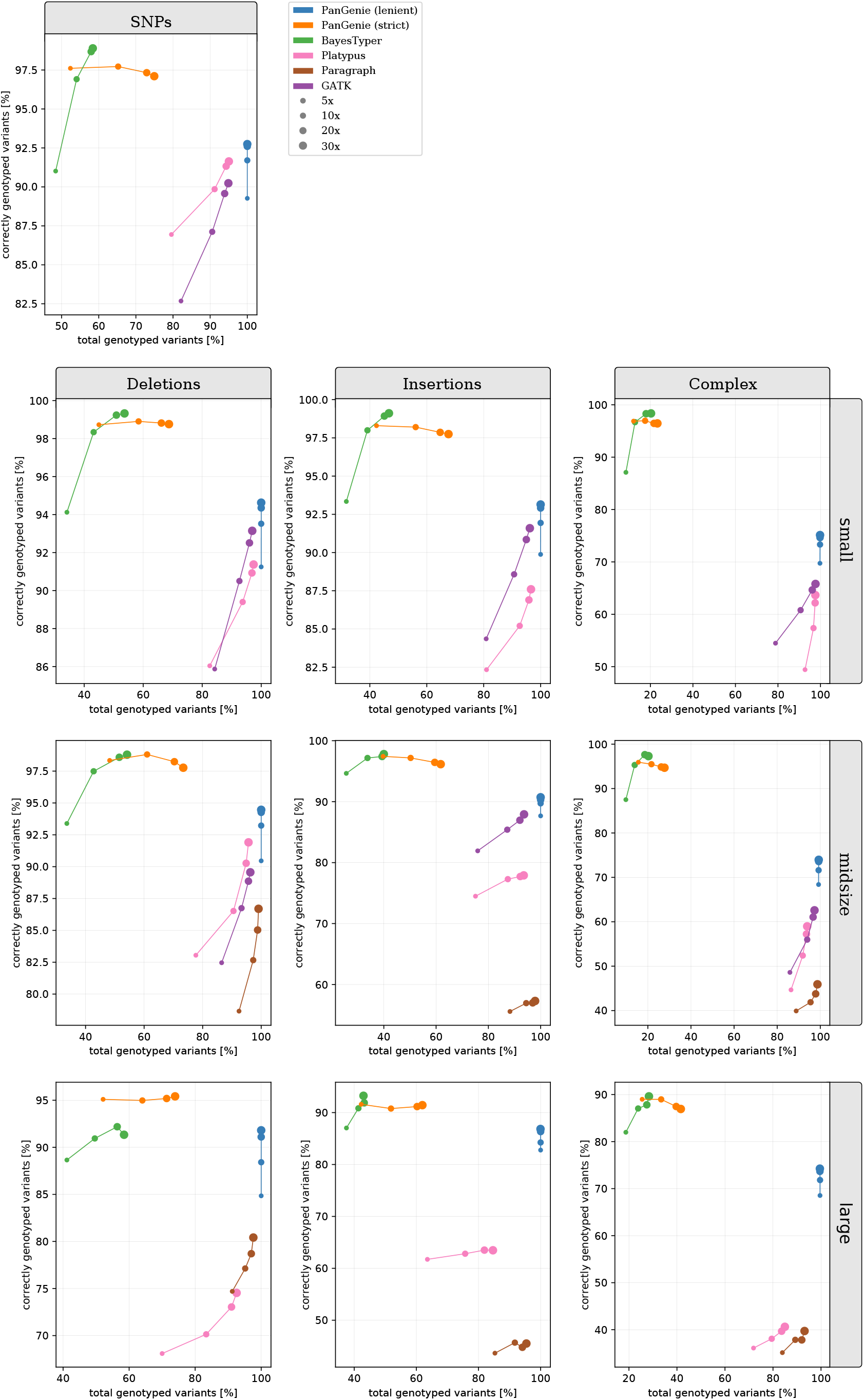
Repeat regions. Genotype performance for sample **HG00731** at different coverages inside of STR/VNTR regions. We ran PanGenie using the **extended** panel, BayesTyper, Paragraph, Platypus and GATK in order to regenotype all callset variants. Besides not applying any filter on the reported genotype qualities (“lenient”), we additionally report genotyping statistics for PanGenie when using “strict” filtering (genotype quality >= 200). Note that GATK was not run on large variants, and Paragraph was only run on midsize and large variants.

For SNPs, all methods reach similar levels of genotype concordances. Platypus and PanGenie (small + extended panel) perform best on the lowest tested coverage of 5×. While PanGenie is able to genotype almost all variants (> 99.998%) using “lenient” filtering on high and low coverage, the other methods show larger levels of variants that they leave untyped. This is especially the case for BayesTyper, which reaches higher levels of genotype concordances than the other tools at coverages 10 − 30− ×, but does not genotype 9% of the SNPs outside of STR/VNTR regions, and 40% inside these repeat regions at coverage 30× for sample HG00731.

For the small variants, PanGenie outperforms the mapping-based approaches on most coverages using the small panel. With the extended panel, we can improve the performance of our method even further, especially inside of STR/VNTR regions. Here, PanGenie reaches genotype concordances superior by 6.5%, 6.26% and 28% compared to the best performing mapping-based approach on insertions, deletions and complex variants, respectively, when using the “lenient” model. BayesTyper produces higher percentages of correct predictions, but is not able to determine genotypes for 30 − 90% of the variants outside of repeat regions, and between 50 − 91% of variants located inside of STR/VNTR regions. Using “strict” filtering, PanGenie is able to reach genotype concordances similar to BayesTyper, while still being able to type much larger fractions of variants.

We observe a similar trend for midsize and large variants as well. Here, PanGenie clearly outperforms the mapping-based tools even when using the small panel of haplotypes. Improvements were largest for large variants inside of repeat regions, where PanGenie with the extended panel and “lenient” filtering is able to reach genotype concordances that are up to 15%, 37% and 89% higher than those of the best performing mapping-based approach for insertions, deletions and complex variants, respectively. The percentages of large variants that could not be genotyped by BayesTyper is between 60 − 80% in all cases, while PanGenie types more than 99% of the variants in each category in “lenient” mode.

When restricting the evaluation to variants contained in the Genome in a Bottle (GIAB) small variant calls [43], PanGenie showed genotyping performances similar to the other methods, while outperforming them on the lowest tested coverage of 5× (Supplementary Section 4.3).

In general, genotyping longer variants based on short-read data is a challenging task, since such variants are often located in repetitive or duplicated regions of the genome [26]. Their short length makes it difficult to unambiguously map the reads in these regions which also effects the genotyping process that relies on these alignments. K-mer based approaches additionally lack the connectivity information contained in the reads, which makes genotyping variants in such difficult regions even more complicated. This is one possible explanation why we observed such high numbers of untyped variants for BayesTyper. PanGenie overcomes these limitations of short reads, as it additionally incorporates long-range haplotype information inherent to the pangenome reference panel it uses. This enables imputation of genotypes in regions poorly covered by k-mers and helps to improve genotyping performance over the other methods, especially for midsize and large variants located in repetitive regions of the genome.

### Runtimes

Runtimes (in CPU hhh:mm:ss) of all methods for sample HG00731 are shown in Table 1. The runtimes for the other samples were very similar and are provided in Supplementary Table 4. For each method, we measured the time required to produce genotypes given an input set of variants and raw, unaligned sequencing reads. Since k-mer based methods PanGenie and BayesTyper bypass the time-consuming read alignment step, they are much faster compared to the remaining, mapping-based methods. GATK and Paragraph were the slowest methods, although they were – unlike the other tools – only run on a subset of variants. PanGenie in contrast, was the fastest method on all coverages. Using the small panel, it was between 3.3 − 3.56× faster than Platypus on the lowest tested coverage of 5×, and between 6.52 − 8.21× faster on coverage 30×. Using the extended panel, these numbers were 1.07 − 1.25× and 4.03 − 4.27×, respectively.

**Table 1:**
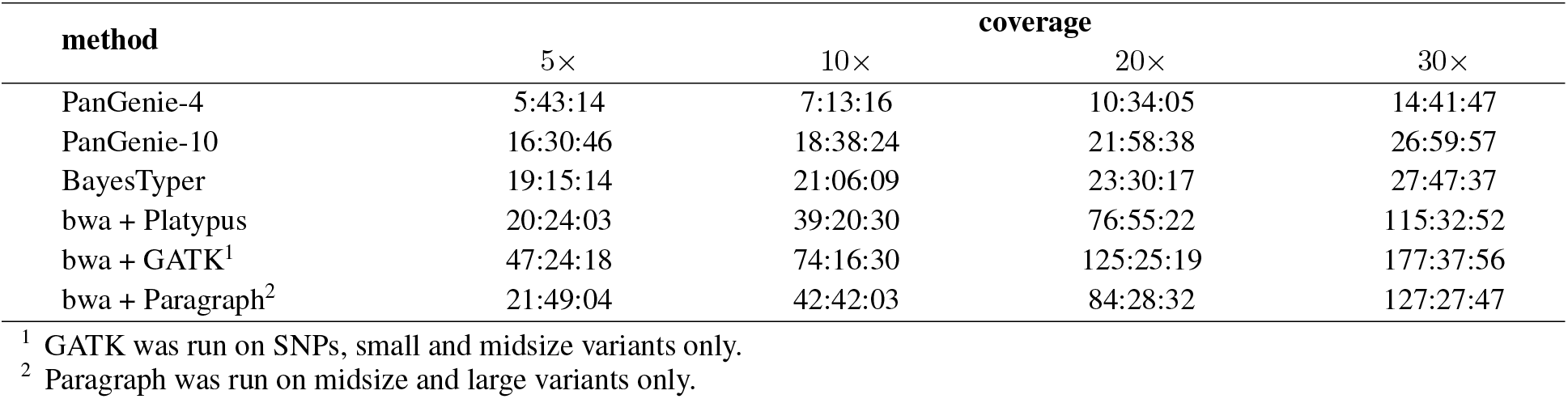
**Runtimes** (in CPU hhh:mm:ss) for sample **HG00731** on all coverage levels. Note that GATK was only run on SNPs, small and midsize variants. Paragraph was only run on midsize and large variants. All other methods were run on all variant types.

| method | coverage |  |  |  |
| --- | --- | --- | --- | --- |
| | $5\times$ | $10\times$ | $20\times$ | $30\times$ |
| PanGenie-4 | 5:43:14 | 7:13:16 | 10:34:05 | 14:41:47 |
| PanGenie-10 | 16:30:46 | 18:38:24 | 21:58:38 | 26:59:57 |
| BayesTyper | 19:15:14 | 21:06:09 | 23:30:17 | 27:47:37 |
| bwa + Platypus | 20:24:03 | 39:20:30 | 76:55:22 | 115:32:52 |
| bwa + GATK<sup>1</sup> | 47:24:18 | 74:16:30 | 125:25:19 | 177:37:56 |
| bwa + Paragraph<sup>2</sup> | 21:49:04 | 42:42:03 | 84:28:32 | 127:27:47 |

<sup>1</sup> GATK was run on SNPs, small and midsize variants only.
<sup>2</sup> Paragraph was run on midsize and large variants only.

#### 2.4 Genotyping larger cohorts

The low runtime of PanGenie makes it well suited to genotype large cohorts. To demonstrate this use case, we applied our tool to a set of 100 randomly selected 1000 Genomes samples based on 1000 Genomes Project high-coverage data (Mike Zody, personal communication). We genotyped all variants contained in the callset that we described in section 2.2 and the 2 × 11 haplotypes contained in our extended panel. We used VCFTools [44] to test the genotype predictions of bi-allelic variants for conformance with Hardy-Weinberg equilibrium and corrected for multiple hypothesis testing by applying Benjamini-Hochberg correction [45] (*α* = 0.05). We skipped such variant positions at which there was a missing genotype for more than 10 samples.

We observed no significant deviation from Hardy-Weinberg equilibrium for 95.7% of all bi-allelic variants. When looking at the different variant types individually, this percentage is between 93.7% and 95.7% (Figure 6), indicating that the genotype predictions made by PanGenie are of good quality. Even for larger structural variants, allele frequencies obtained from our variant predictions largely behave as expected by Hardy-Weinberg equilibrium (Figure 6). At the same time, PanGenie on average only took about 30 CPU hours per sample, demonstrating the scalability of our tool.

**Figure 6:**
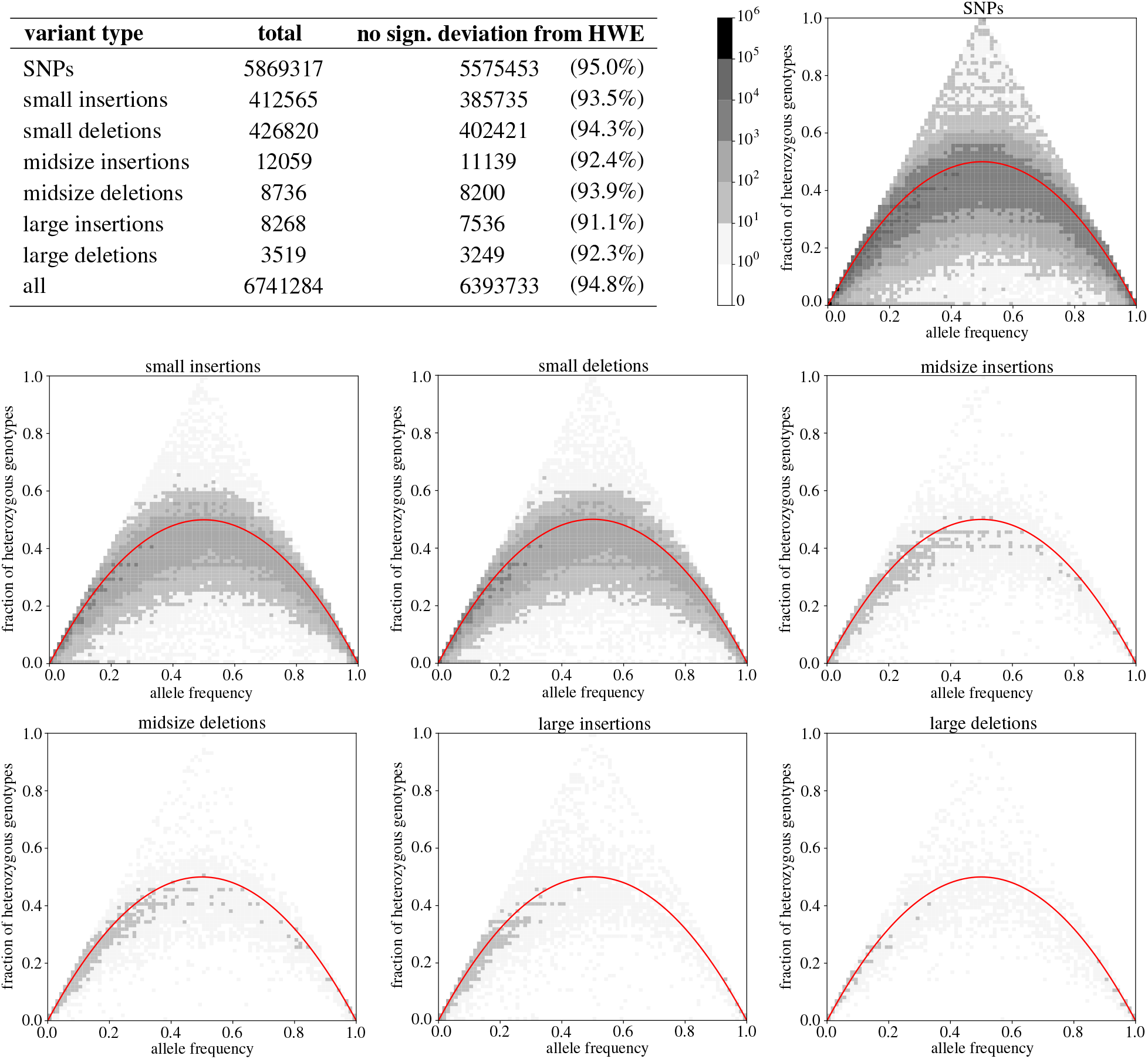
Genotyping larger cohorts. The table provides the amount of variants for which no significant deviation from Hardy-Weinberg Equilibrium was observed. Each plot shows the fraction of heterozygous genotypes at a variant position as a function of the allele frequency. The red curve shows what is expected according to Hardy-Weinberg equilibrium.

## 3 Discussion

We presented an algorithm which uses k-mer counts from short read sequencing data together with a panel of haplotype-resolved assemblies to genotype a yet uncharacterized sample. We show how to formulate this problem in terms of a Hidden Markov Model that models each of the haplotypes of the sample in question as a mosaic of the given haplotype sequences. This algorithm is fast since it bypasses the expensive read alignment step and can also genotype variants located in repetitive or duplicated regions of the genome that are usually poorly covered by unique k-mers. We believe that this is the first approach which can leverage the long-range haplotype information inherent to a panel of assembled haplotype sequences in combination with read k-mer counts for genotyping a new sample. While we generated such pangenome reference panels from haplotype-resolved assemblies for this work, we want to stress that generating these panels was not the main focus of this paper and that our genotyping algorithm is not restricted to panels created in this way. In fact, it can be applied to any acyclic pangenome graph which is represented as a fully-phased, multisample VCF file.

Our experiments showed that PanGenie works as well as mapping-based approaches for small variants, and at the same time, was able to genotype larger fractions of variants compared to the other k-mer based method BayesTyper. Especially for large and midsize variants, PanGenie clearly outperforms mapping-based approaches, while again, compared to BayesTyper, being able to provide genotypes for much larger amounts of variants not typable by the latter. At the same time, our approach was faster than the other methods, especially when comparing to the mapping-based approaches which require alignments of reads to a reference genome. The fast runtime of our method also makes it well suited for genotyping larger cohorts.

We hence have presented a method that is both scalable and leverages a haplotype-resolved pangenome reference to enable genotyping of otherwise inaccessible variants. Still, some limitations remain. Since we assume that the unknown haplotypes of the sample to be genotyped are mosaics of the given panel haplotypes, it currently cannot be used in order to genotype rare variants that are only present in the sample, but in none of the other haplotypes. Here, we believe that there are exciting opportunities to define downstream workflow that only discover variation that our approach has not captured because it was not present in the reference panel. That is, one could filter the reads for yet “unexplained” k-mers and use those for the discovery of rare variants.

The runtime of our method depends on the number of input haplotypes, as we define a hidden state for each possible pair of haplotypes that can be selected at a variant position. Therefore, additional engineering will be required to use much larger panels, which could be approached similarly to how statistical phasing packages prune the solution space and/or proceed iteratively.

All in all, we have presented a method that we see as a way forward once high-quality phased reference assemblies become widely available to the genomics community while, at the same time, the size of disease cohort used in association studies grows further.

## 4 Methods

The input to our genotyping algorithm is a reference genome (FASTA-file), short-read sequencing reads (FASTQ format) and a multisample VCF file that defines a pangenome graph containing variants and known haplotype sequences. In order to create such an input VCF, we have developed a pipeline which calls variants from haplotype resolved assemblies as described below. However, we want to stress that our tool is not restricted to VCFs created in this way and in fact can be run with any fully phased, multisample VCF file.

### 4.1 Pangenome reference construction

We used haplotype-resolved assemblies of five individuals (HG00731, HG00732, NA12878, NA24385 and PGP1) [34, 35] and separately aligned the contigs of each haplotype to the reference genome (GRCh38). This was done using minimap2 [46] with parameters `-cx asm5 –-cs`. Next, we called variants on each haplotype using paftools (https://github.com/lh3/minimap2/tree/master/misc) with default parameters. We only kept variants located in regions in which all haplotypes were covered by exactly one contig alignment in order to filter out low quality or erroneous calls. All other regions, in which at least one of the haplotypes was covered by none or multiple contig alignments, were excluded from further analyses.

Our goal is to construct an acyclic and directed graph by inserting the variants of all haplotypes into the linear reference genome. Each variant produces a bubble in the graph whose branches define the corresponding alleles. The input haplotypes can be represented as paths through the resulting pangenome. When constructing the graph, we represent sets of variants overlapping across haplotypes as a single bubble with potentially multiple branches reflecting all the allele sequences observed in the haplotypes in the respective genomic region. See Figure 2 for an example. We represent the pangenome in terms of a fully phased, multi-sample VCF file that contains an entry for each bubble in the graph. At each site, the number of alternative alleles is limited by the number of input haplotype sequences and the genotypes of each sample define two paths through this graph corresponding to the respective haplotypes.

We extended the number of haplotype paths in the graph by using PanGenie to phase additional samples based on short read sequencing data and the paths already present in our graph. This is achieved by applying the Viterbi algorithm to our Hidden Markov Model (see Section 4.3 for details). In this way, we added haplotypes of six additional individuals to the graph. These include samples of Chinese and Yorubian descent (HG00512, NA19238) as well as four samples from different populations (see Figure 3a) The underlying reads for the Chinese and Yorubian samples were obtained from [3] and those of the remaining samples from 1000 Genomes Project high-coverage data (Mike Zody, personal communication). We used bcftools (https://github.com/samtools/bcftools), VCFTools [44] and `vcfstats` from Real Time Genomics [47] to generate the callset statistics presented in Figure 3.

The individuals of Puerto Ricean, Chinese and Yorubian descent, are part of three trios. We additionally determined the genotypes for the remaining samples (HG00733, HG00513 and HG00514, NA19239 and NA19240). This was done in a similar way as for the other samples, using haplotype-resolved assemblies to determine phasings for HG00733, and short-read sequencing reads in order to phase the remaining samples, for which such assemblies were not available to us. Using the trio information, we can check whether the variant calls are consistent with the laws of Mendelian inheritance. For the Puerto Rican trio, we observed 98.34% Mendelian consistent genotypes for the phasings produced from haplotype-resolved assemblies. For the Chinese and Yorubian trios, these percentages were 98.76% and 97.96%, respectively, for the phasings produced by PanGenie. For further analysis, we removed all variants from our graph for which there was a Mendelian error in at least one of the trios.

### 4.2 Identifying unique k-mers

Sets of variants that are less than the k-mer size apart (we use *k* = 31) are combined and treated as a single variant position. The alleles corresponding to such a combined variant are defined by the haplotype paths in the respective region. For each variant position *v*, we determine a set of k-mers, *kmers*_*v*_, that uniquely characterize the variant region. This is done by counting all k-mers along haplotype paths in the pangenome graph using Jellyfish [36], and then determining a set of k-mers for each variant, that occur at most once within a single allele sequence and are not found anywhere outside of the variant bubble. We additionally count all k-mers of the graph in the sequencing reads.

### 4.3 Hidden Markov Model

We define a Hidden Markov Model that can be used to compute the two most likely haplotype sequences of a given sample based on known haplotype paths and the sample reads. The new haplotype sequences are combinations of the existing paths through the graph and are computed based on the copy numbers of unique k-mers observed in the sequencing reads provided for the sample to be genotyped.

#### Hidden States and Transitions

We assume to be given *N* haplotype paths *H*_*i*_, *i* = 1, …, *N*, through the graph. Furthermore, for each variant position *v, v* = 1, …, *M*, we are given a vector of k-mers, *kmers*_*v*_ that uniquely characterize the alleles of a variant. We assume some (arbitrary) order of the elements in *kmers*_*v*_ and refer to the *i*th k-mer as *kmers*_*v*_[*i*]. Additionally, we are given sequencing data of the sample to be genotyped and corresponding k-mer counts for all k-mers in *kmers*_*v*_. For each variant position *v*, we define set of hidden states ℋ_*v*_ = {*H*_*v,i,j*_ | ≤ *N*} which contains a state for each possible pair of the *N* given haplotype paths in the graph. Each such state *H*_*v,i,j*_ induces an assignment of copy numbers to all k-mers in *kmers*_*v*_ defined as shown below.

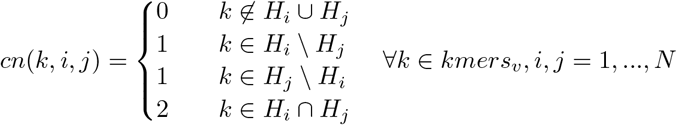

The idea here is that we expect to see copy number 2 for all k-mers occurring on both haplotype paths. In case only one of the haplotypes contains a k-mer, its copy number must be 1 and k-mers that do not appear in any of the two paths must have copy number 0. Thus, for each state *H*_*v,i,j*_ in ℋ_*v*_, we define the vector *a*_*v,i,j*_ that contains the assigned copy numbers for all k-mers, i.e. *a*_*v,i,j*_[*l*] = *cn*(*kmers*_*v*_[*l*], *i, j*).

From each state *H*_*v,i,j*_ ∈ ℋ_*v*_ that corresponds to variant position *v*, there is a transition to each state corresponding to the next position, *v* + 1. Additionally, there is a *start* state, from which there is a transition to each state of the first variant, and an *end* state, to which there is a transition from each state that corresponds to the last variant position. See Figure 1c for an example.

#### Transition Probabilities

Transition probabilities are computed similar to how the Li-Stephans model [28] defines them. We assume to be given a recombination rate *r* and the effective population size *N*_*e*_. For two ascending variant positions *v* − 1 and *v* that are *x* bases apart in the genome, we first compute the genetic distance:

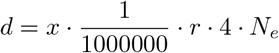

We further compute the Li-Stephans transition probabilities as:

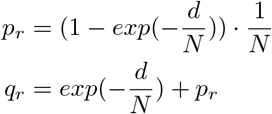

Finally, the transition probability from state *H*_*v*− 1,*i,j*_ to state *H*_*v,k,l*_ is computed as shown below.

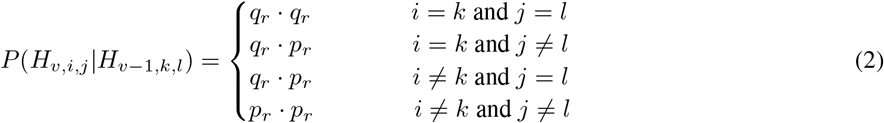

#### Observable States

Each hidden state *H*_*v,i,j*_ ∈ *ℋ*_*v*_ outputs a count for each k-mer in *kmers*_*v*_. Let *obs*(*k*) be a function that returns the observed count in the reads of a k-mer *k* ∈ *𝒦* and the vector such that 𝒪_*v*_[*l*] = *obs*(*kmers*_*v*_[*j*]). In order to define the emission probabilities, we first need to model the distribution of k-mer counts for each copy number, *P* (*obs*(*k*) *cn*(*k*) = *i*), *i* = 0, 1, 2. For copy number 2, we use a Poission distribution whose mean *λ* we compute from the read k-mer-count histogram. Similarly, we approximate the k-mer count distribution for copy number 1 in terms of a Poisson distrubution with mean *λ/*2. For copy number 0, we need to model the erroneous k-mers that arise from sequencing errors. This is done using a Geometric distribution. whose parameter *p* we choose based on the mean k-mer coverage. Finally, we compute the emission probability for a given state and given observed read k-mer counts as shown below, making the assumption that the k-mer counts are independent.

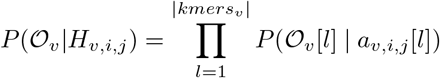

#### Genotypes and Haplotypes

In this model, genotypes correspond to pairs of given haplotype paths at each variant position. Genotype likelihoods can be computed using the Forward-Backward algorithm, and haplotype sequences can be computed by running Viterbi. We assume to have observed copy number *obs*(*k*) of each unique k-mers in 𝒦.

#### Forward-Backward algorithm

The initial distribution of our HMM is such that we assign probability 1 in the *start* state and 0 to all others. Forward probabilities *α*_*v*_() are computed in the following way.

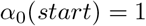

For states corresponding to variant position *v* = 1, …, *M*, the Forward probabilities are computed as shown below. The set of observed k-mer counts at position *v* is given by *𝒪*_*v*_ = {*obs*(*k*), *k* ∈ *kmers*_*v*_}.

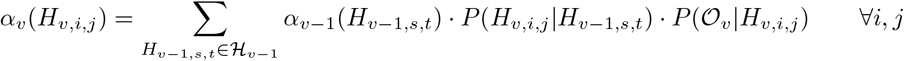

The transition probabilities are computed as described above, except for transitions from the *start* state to all states in the first column, which we assume to have uniform probabilities.

Backward probabilities are computed in a similar manner. We set

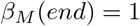

For *v* = 1, …, *M* − 1, we compute them as

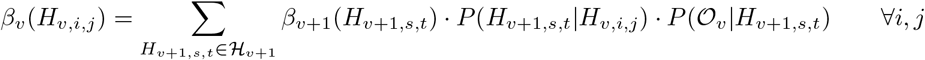

Finally, posterior probabilities for the states can be computed.

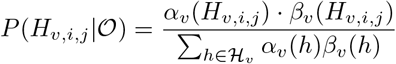

Several states at a variant position *v* can correspond to the same genotype, as different paths can cover the same allele. Also, the alleles in a genotype are unordered, therefore states *H*_*v,i,j*_ and *H*_*v,j,i*_ always lead to the same genotype. In order to compute genotype likelihoods, we sum up the posterior probabilities for all states that correspond to the same genotype. In this way, we can finally compute genotype likelihoods for all genotypes at a variant position, based on which a genotype prediction can be made.

#### Viterbi algorithm

In order to get the haplotype sequences, we can compute the two haplotypes underlying the Viterbi path. We again start in the *start* state.

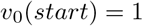

For the other positions *v* = 1, …, *M*, we compute:

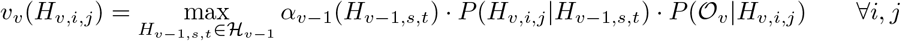

We finally obtain the Viterbi path by backtracking.

## Availability of data and materials

Code to reproduce the data and rerun the analysis is available at https://bitbucket.org/jana_ebler/genotyping-experiments/src/master/.

The implementation of PanGenie is available at https://bitbucket.org/jana_ebler/pangenie/src/master/.

## Supplementary material

### Callset statistics

Table 2 shows the numbers of variants of each type that were present in the callset constructed from reference-resolved assemblies. For each sample, we show the total number of variants present in at least one of its haplotypes, i.e. all variants for which the sample has a genotype different from 0/0 (total), as well as the number of variants for which a sample carried at least one allele not seen in any of the remaining samples (unique). All variants that are unique to a sample will not be genotypable by our HMM based approach, since the assumption underlying our model is that the unknown haplotypes can be constructed as a mosaic of the haplotypes already known. Thus, if the sample in question carries an allele not seen before, it cannot be correctly genotyped with such a re-typing approach.

**Table 2:**
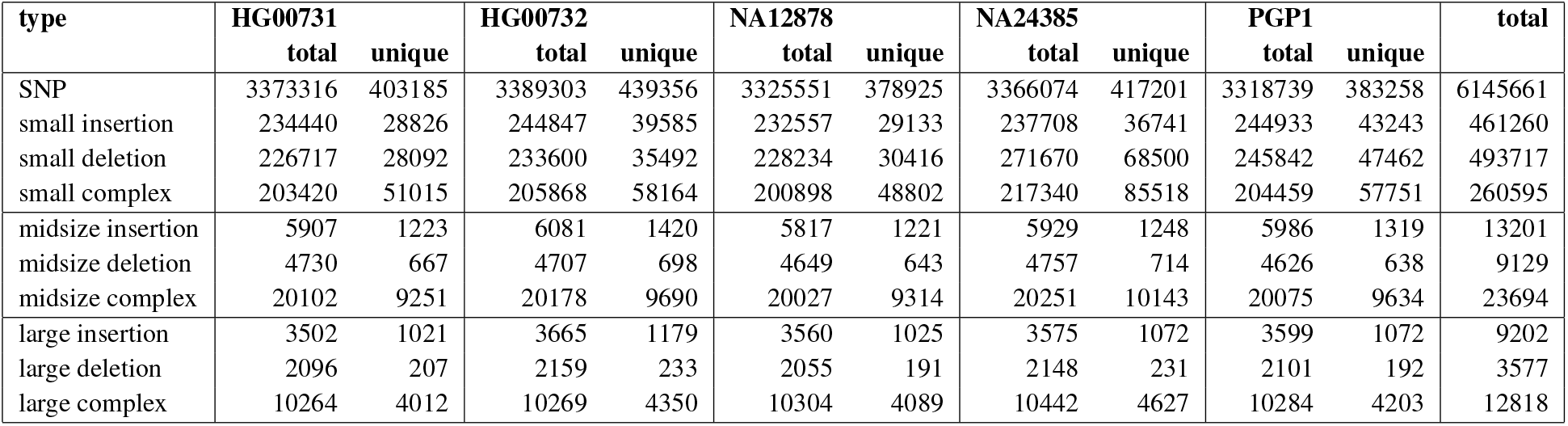
Variant statistics. Total number of variants detected in each sample, as well as the number of variants for which a sample carried an allele not present in the other samples.

| type | HG00731 |  | HG00732 |  | NA12878 |  | NA24385 |  | PGP1 |  | total |
| --- | --- | --- | --- | --- | --- | --- | --- | --- | --- | --- | --- |
|  | total | unique | total | unique | total | unique | total | unique | total | unique |  |
| SNP | 3373316 | 403185 | 3389303 | 439356 | 3325551 | 378925 | 3366074 | 417201 | 3318739 | 383258 | 6145661 |
| small insertion | 234440 | 28826 | 244847 | 39585 | 232557 | 29133 | 237708 | 36741 | 244933 | 43243 | 461260 |
| small deletion | 226717 | 28092 | 233600 | 35492 | 228234 | 30416 | 271670 | 68500 | 245842 | 47462 | 493717 |
| small complex | 203420 | 51015 | 205868 | 58164 | 200898 | 48802 | 217340 | 85518 | 204459 | 57751 | 260595 |
| midsize insertion | 5907 | 1223 | 6081 | 1420 | 5817 | 1221 | 5929 | 1248 | 5986 | 1319 | 13201 |
| midsize deletion | 4730 | 667 | 4707 | 698 | 4649 | 643 | 4757 | 714 | 4626 | 638 | 9129 |
| midsize complex | 20102 | 9251 | 20178 | 9690 | 20027 | 9314 | 20251 | 10143 | 20075 | 9634 | 23694 |
| large insertion | 3502 | 1021 | 3665 | 1179 | 3560 | 1025 | 3575 | 1072 | 3599 | 1072 | 9202 |
| large deletion | 2096 | 207 | 2159 | 233 | 2055 | 191 | 2148 | 231 | 2101 | 192 | 3577 |
| large complex | 10264 | 4012 | 10269 | 4350 | 10304 | 4089 | 10442 | 4627 | 10284 | 4203 | 12818 |

### Comparison to gnomAD

We compared the variant calls that we obtained from haplotype-resolved assemblies of five individuals to the variants that are part of the Genome Aggregation Database (gnomAD) [4]. gnomAD contains 433,371 structural variants collected across 14,891 genomes from different populations. Since gnomaD calls were generated relative to reference genome version hg19, we used UCSC liftOver (https://genome.ucsc.edu/cgi-bin/hgLiftOver) to convert their coordinates to hg38. We compared the variants contained in gnomAD to our assembly-based variant calls. We excluded variants genotyped with an allele frequency of 0.0 across all 100 genotyped samples (Section 2.4). We determined all variants with a reciprocal overlap of at least 50% between the gnomAD calls and our assembly-based callset (chromosomes 1-22) and found that both callsets had 6,398 variants in common. 368,530 variants were only contained in gnomAD and 21,370 were only in our assembly callset. 35.3% of the 6,398 variants in the intersection are located inside of STR/VNTR regions. For the variants contained only in our assembly callset, this percentage is around 80%. We suspect that the reason these variants cannot be found in gnomAD might be that such repetitive regions are not accessible by short read data used to produce the gnomAD variant calls. For each variant in our assembly callset, we further computed the distance to the closest gnomAD variant. Additionally, we used `bedtools shuffle` [48] to randomly permute the variants among the genome. Then we again determined distances to the closest gnomAD variants. We show the resulting distances in Figure 7.

**Figure 7:**
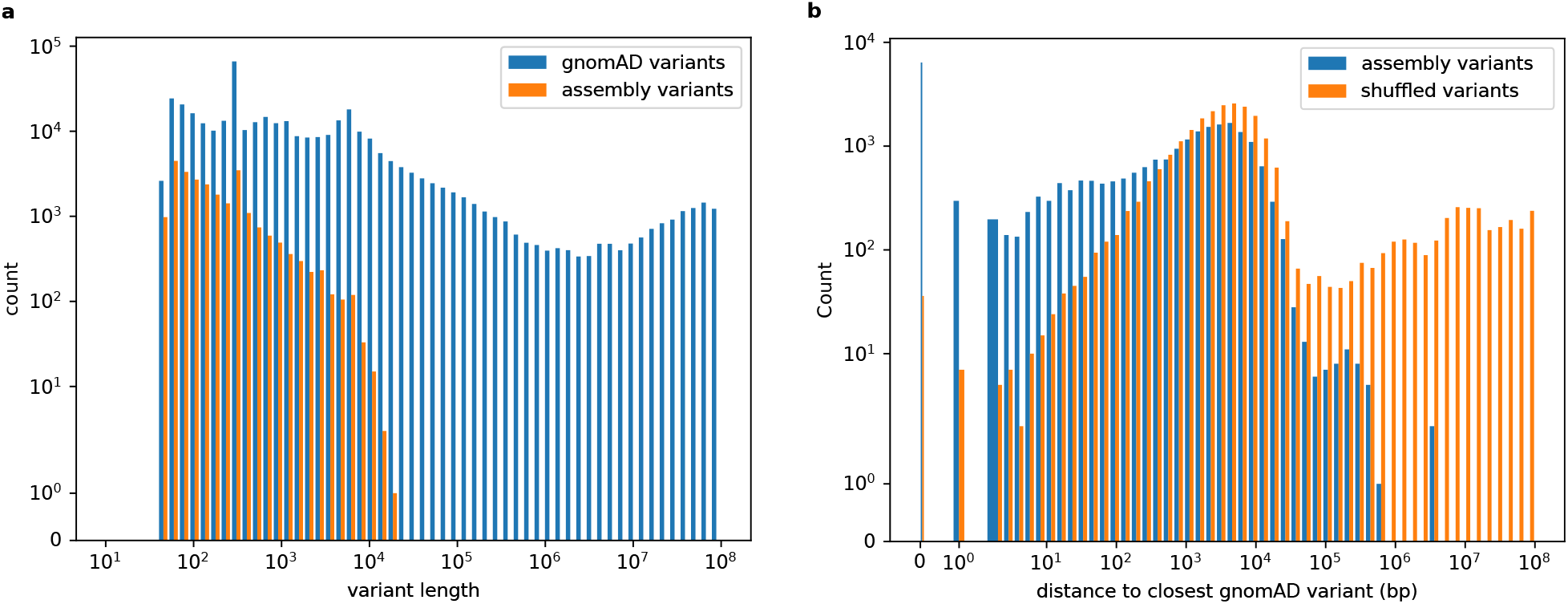
Comparison to gnomAD. **a)** Histogram of the variant length of the structural variants contained in the gnomAD callset (blue) and our assembly callset (orange). **b)** For each variant in our assembly callset, we computed the distance to the closest gnomAD variant (blue). We repeated the same analysis after randomly permuting our variant calls along the reference genome (orange).

### Benchmarking results

#### STR/VNTR regions

Especially structural variants tend to be located in repetitive and more complicated regions of the genome. For all variants that we genotyped in sample HG00731 (Section 2.3), we show the number of sites located inside and outside of STR/VNTR regions which we obtained from the UCSC genome browser [42]. The numbers are presented in Table 3. It can be observed that the majority (between 54 − 79%) of midsize and large variants are indeed inside of repetitive regions.

**Table 3:** Repetitive regions. Shown are the number and percentages of variants located inside and outside of STR/VNTR regions for sample HG00731.

| variant type | all regions | non-repetitive regions |  | repeat regions |  |
| --- | --- | --- | --- | --- | --- |
| SNP | 5742475 | 5527879 | 96,26% | 214596 | 3,74% |
| small deletion | 465625 | 401391 | 86,20% | 64234 | 13,80% |
| small insertion | 432434 | 382244 | 88,39% | 50190 | 11,61% |
| small complex | 209580 | 144809 | 69,09% | 64771 | 30,91% |
| midsize deletion | 8462 | 2971 | 35,11% | 5491 | 64,89% |
| midsize insertion | 11978 | 5497 | 45,89% | 6481 | 54,11% |
| midsize complex | 14443 | 3092 | 21,41% | 11351 | 78,59% |
| large deletion | 3370 | 1100 | 32,64% | 2270 | 67,36% |
| large insertion | 8178 | 3220 | 39,37% | 4958 | 60,63% |
| large complex | 8806 | 2071 | 23,52% | 6735 | 76,48% |

**Table 4:** **Runtimes** (in CPU hhh:mm:ss) of the different genotyping methods at different coverages.

| coverage | method | runtime (CPU sec) |  |  |  |
| --- | --- | --- | --- | --- | --- |
|  |  | HG00731 | HG00732 | NA12878 | NA24385 |
| 5 | PanGenie-4 | 5:43:14 | 5:47:27 | 5:24:09 | 5:50:16 |
|  | PanGenie-10 | 16:30:46 | 16:09:53 | 16:33:56 | 16:39:10 |
|  | BayesTyper | 19:15:14 | 19:01:26 | 19:15:50 | 19:18:32 |
|  | bwa + Platypus | 20:24:03 | 20:17:03 | 17:49:02 | 20:23:04 |
|  | bwa + GATK<sup>1</sup> | 47:24:18 | 46:41:48 | 43:53:18 | 46:28:50 |
|  | bwa + Paragraph<sup>2</sup> | 21:49:04 | 21:42:20 | 19:42:04 | 21:58:48 |
| 10 | PanGenie-4 | 7:13:16 | 7:21:35 | 6:53:07 | 7:28:14 |
|  | PanGenie-10 | 18:38:24 | 18:32:37 | 17:58:22 | 18:39:40 |
|  | BayesTyper | 21:06:09 | 20:48:47 | 20:44:49 | 21:32:58 |
|  | bwa + Platypus | 39:20:30 | 39:00:07 | 33:41:37 | 39:29:54 |
|  | bwa + GATK<sup>1</sup> | 74:16:30 | 73:38:42 | 67:07:30 | 73:17:21 |
|  | bwa + Paragraph<sup>2</sup> | 42:42:03 | 42:20:31 | 38:30:50 | 43:26:37 |
| 20 | PanGenie-4 | 10:34:05 | 10:17:36 | 9:03:41 | 10:16:48 |
|  | PanGenie-10 | 21:58:38 | 21:12:08 | 19:39:27 | 21:24:34 |
|  | BayesTyper | 23:30:17 | 23:14:27 | 22:28:14 | 24:27:43 |
|  | bwa + Platypus | 76:55:22 | 76:40:07 | 65:51:00 | 77:35:07 |
|  | bwa + GATK<sup>1</sup> | 125:25:19 | 124:35:58 | 113:03:27 | 124:42:22 |
|  | bwa + Paragraph<sup>2</sup> | 84:28:32 | 84:03:30 | 77:15:57 | 86:23:53 |
| 30 | PanGenie-4 | 14:41:47 | 17:34:07 | 12:59:47 | 14:07:18 |
|  | PanGenie-10 | 26:59:57 | 26:38:29 | 23:42:00 | 23:49:38 |
|  | BayesTyper | 27:47:37 | 27:27:24 | 25:18:09 | 30:04:13 |
|  | bwa + Platypus | 115:32:52 | 114:32:36 | 95:31:42 | 115:59:24 |
|  | bwa + GATK<sup>1</sup> | 177:37:56 | 175:12:35 | 152:07:04 | 175:26:41 |
|  | bwa + Paragraph<sup>2</sup> | 127:27:47 | 126:00:29 | 115:11:40 | 129:51:49 |

<sup>1</sup> GATK was run on SNPs, small and midsize variants only.
<sup>2</sup> Paragraph was run on midsize and large variants only.

**Table 5:** GIAB overlap. Number of variants that overlap with the Genome in a Bottle small variant calls.

|  | **SNPs** | **small deletions** | **small insertions** | **midsize deletions** | **midsize insertions** |
| --- | --- | --- | --- | --- | --- |
| **GIAB callset** | 3085616 | 247499 | 242711 | 2658 | 1925 |
| **variants in overlap** | 2534770 | 182493 | 183257 | 1613 | 1390 |
| **overlap [%]** | 82,15% | 73,73% | 75,50% | 60,68% | 72.21% |

#### Results for the extended panel

We additionally show the genotyping results of all methods using the extended panel for samples HG00732, NA12878 and NA24385 in Figures 8-13. Genotyping experiments where run in the same way as for HG00731 presented in Section 2.3. For PanGenie, we used the extended panel that contained 10 samples (20 haplotypes). Besides using all output genotypes produced by PanGenie regardless of the reported genotype quality (“lenient”), we additionally report results of PanGenie when applying a much more strict filtering using genotype quality score of 200 (“strict”). For all other tools, we used all genotypes that they reported and did not use any filtering on genotype qualities. We again show results for variants inside and outside of repetitive regions.

**Figure 8:**
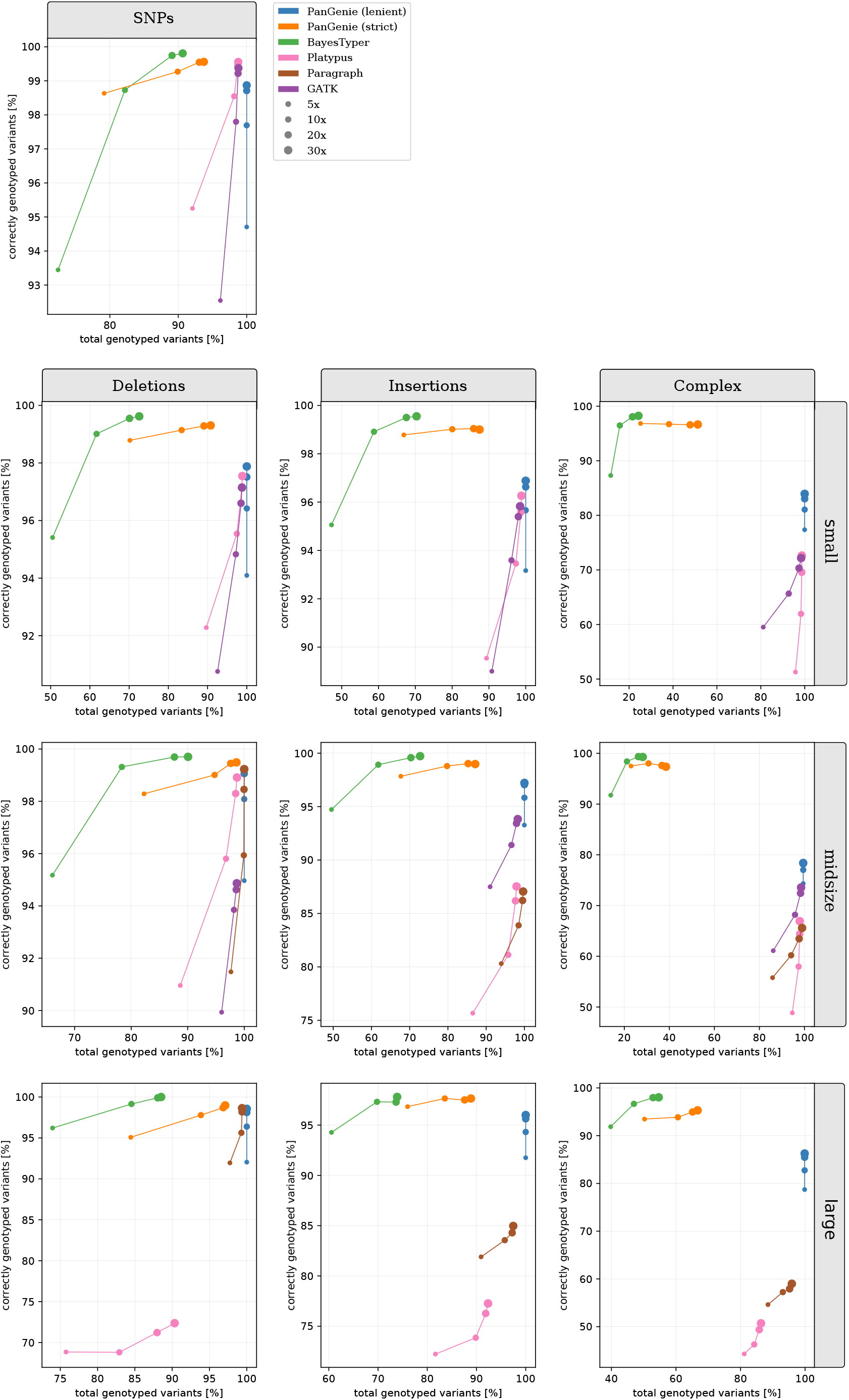
extended panel results for HG00732 in non-repetitive regions. We ran PanGenie using the **extended** panel, BayesTyper, Paragraph, Platypus and GATK in order to re-genotype all callset variants located outside of repetitive STR/VNTR regions. Besides not applying any filter on the reported genotype qualities (“lenient”), we additionally report genotyping statistics for PanGenie when using “strict” filtering (genotype quality >= 200). Note that GATK was not run on large variants, and Paragraph was only run on midsize and large variants.

**Figure 9:**
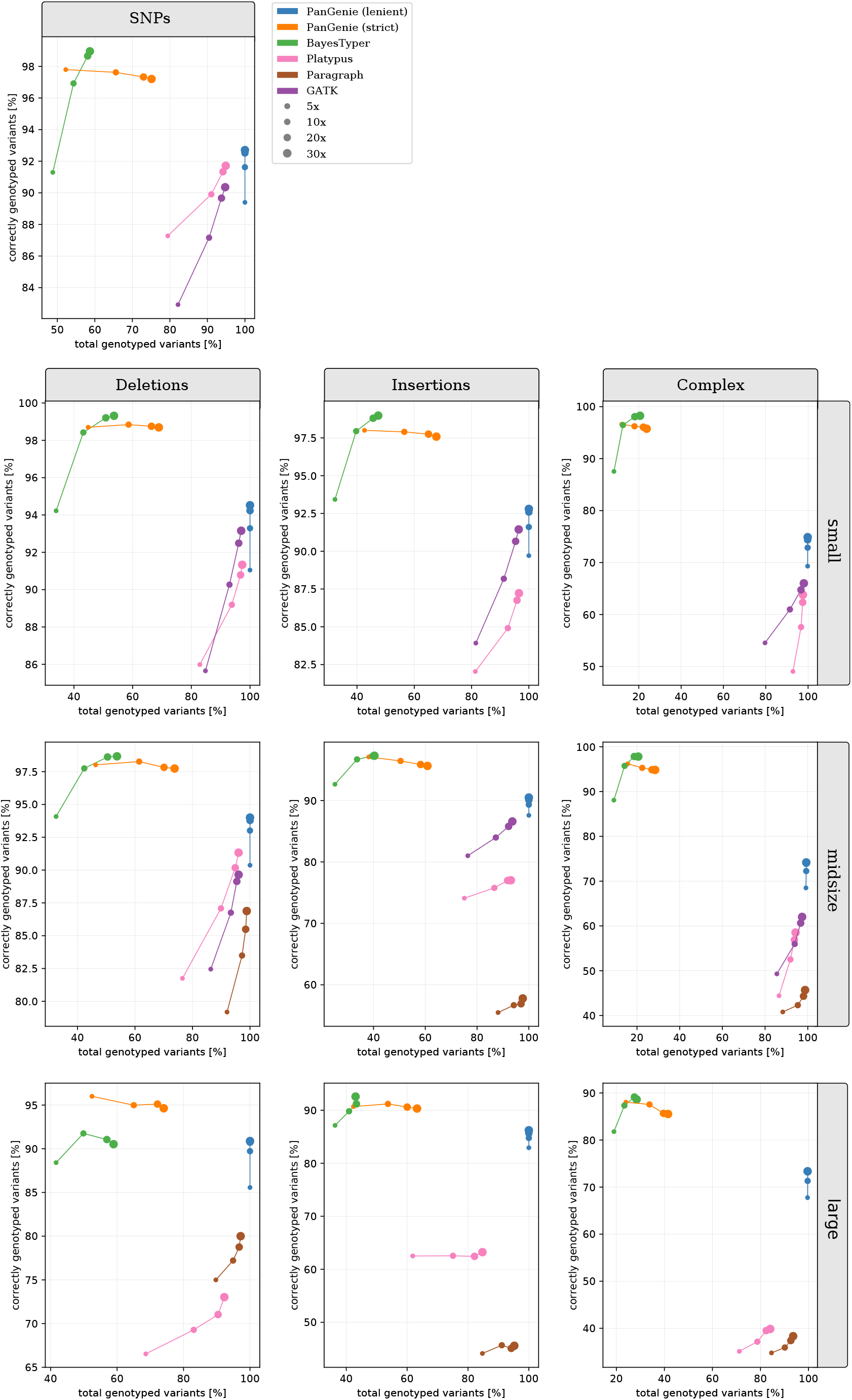
extended panel results for HG00732 in repeat regions. We ran PanGenie using the **extended** panel, BayesTyper, Paragraph, Platypus and GATK in order to re-genotype all callset variants located inside of repetitive STR/VNTR regions. Besides not applying any filter on the reported genotype qualities (“lenient”), we additionally report genotyping statistics for PanGenie when using “strict” filtering (genotype quality >= 200). Note that GATK was not run on large variants, and Paragraph was only run on midsize and large variants.

**Figure 10:**
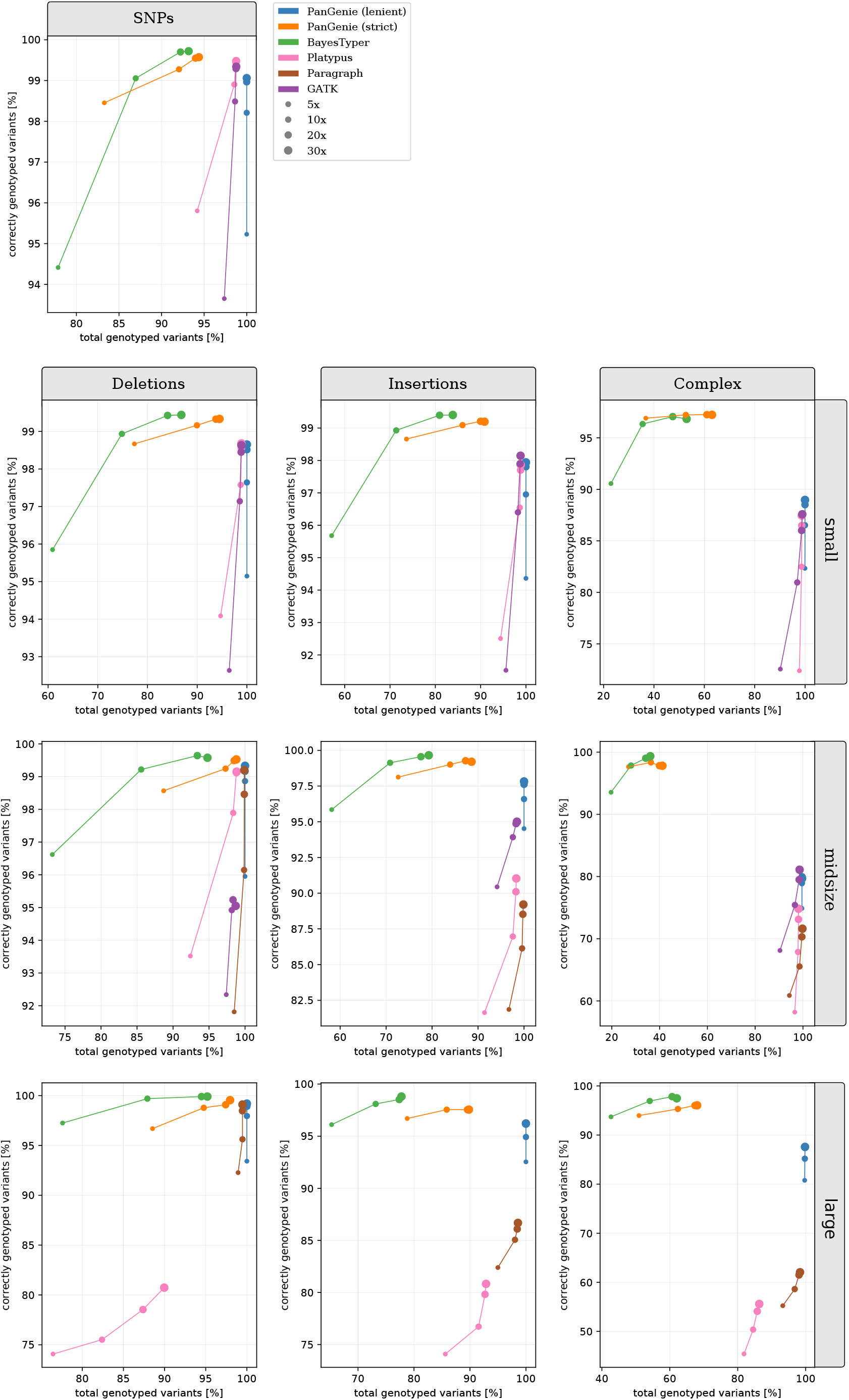
extended panel results for NA12878 in non-repetitive regions. We ran PanGenie using the **extended** panel, BayesTyper, Paragraph, Platypus and GATK in order to re-genotype all callset variants located outside of repetitive STR/VNTR regions. Besides not applying any filter on the reported genotype qualities (“lenient”), we additionally report genotyping statistics for PanGenie when using “strict” filtering (genotype quality >= 200). Note that GATK was not run on large variants, and Paragraph was only run on midsize and large variants.

**Figure 11:**
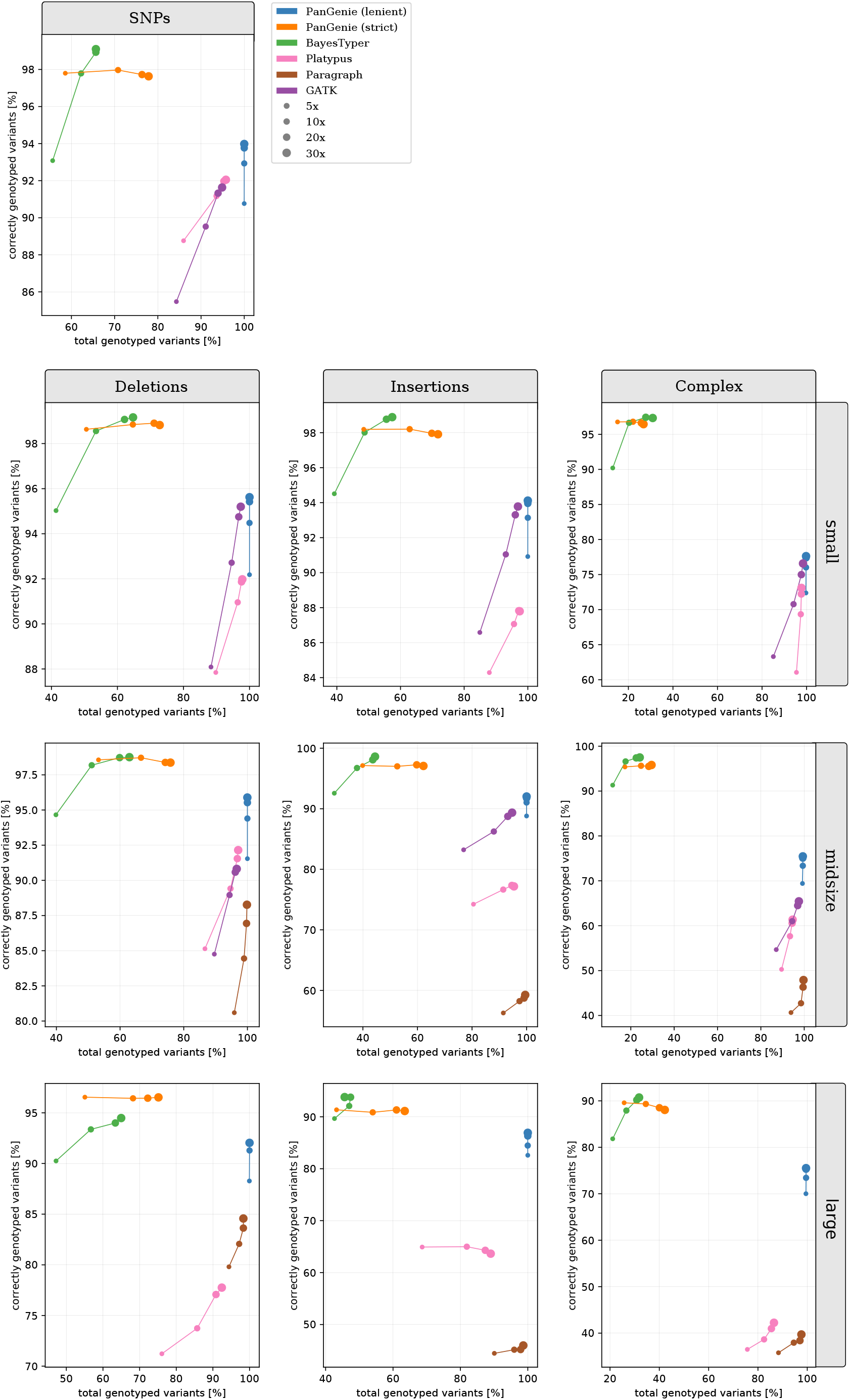
extended panel results for NA12878 in repeat regions. We ran PanGenie using the **extended** panel, BayesTyper, Paragraph, Platypus and GATK in order to re-genotype all callset variants located inside of repetitive STR/VNTR regions. Besides not applying any filter on the reported genotype qualities (“lenient”), we additionally report genotyping statistics for PanGenie when using “strict” filtering (genotype quality >= 200). Note that GATK was not run on large variants, and Paragraph was only run on midsize and large variants.

**Figure 12:**
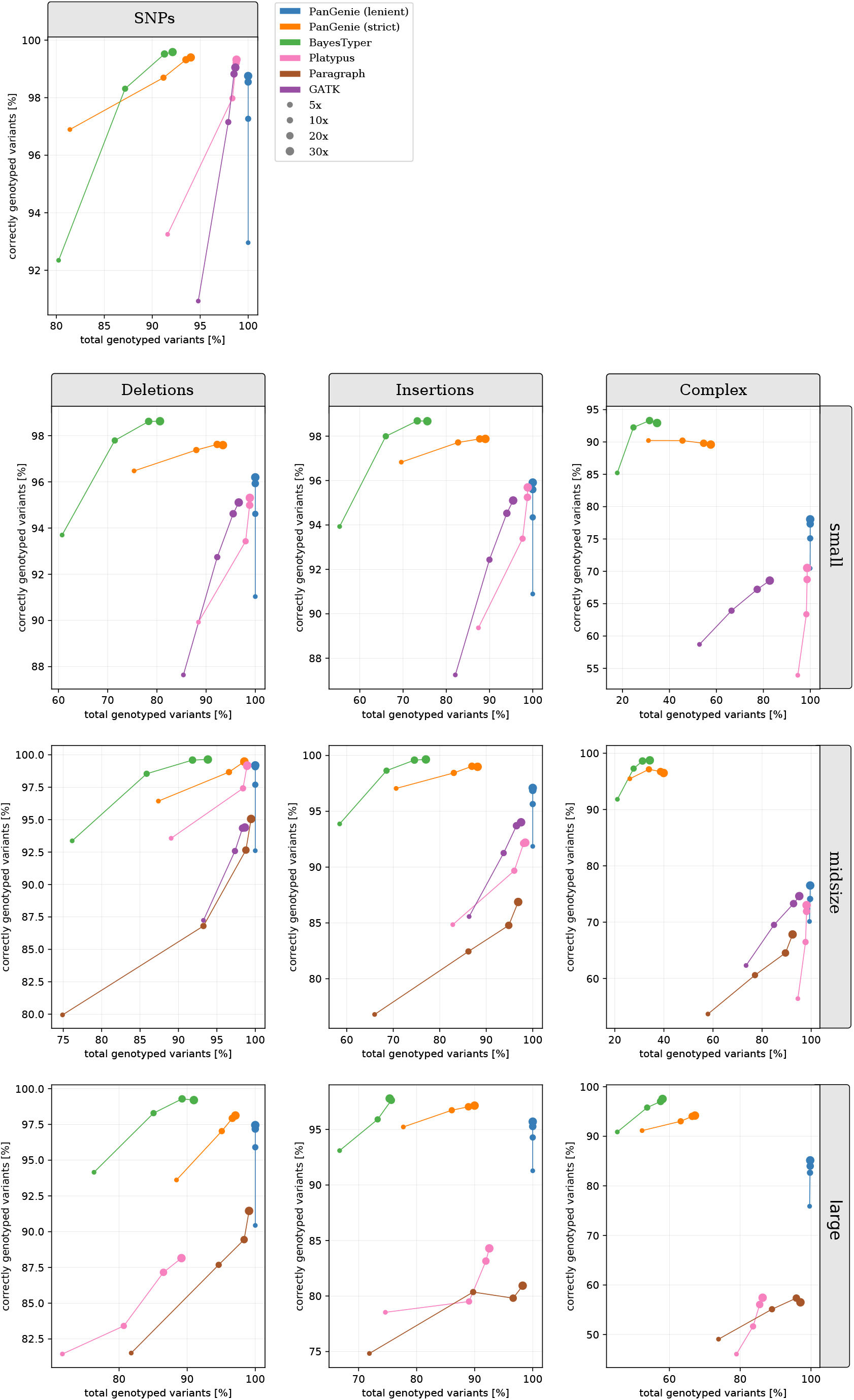
extended panel results for NA24385 in non-repetitive regions. We ran PanGenie using the **extended** panel, BayesTyper, Paragraph, Platypus and GATK in order to re-genotype all callset variants located outside of repetitive STR/VNTR regions. Besides not applying any filter on the reported genotype qualities (“lenient”), we additionally report genotyping statistics for PanGenie when using “strict” filtering (genotype quality >= 200). Note that GATK was not run on large variants, and Paragraph was only run on midsize and large variants.

**Figure 13:**
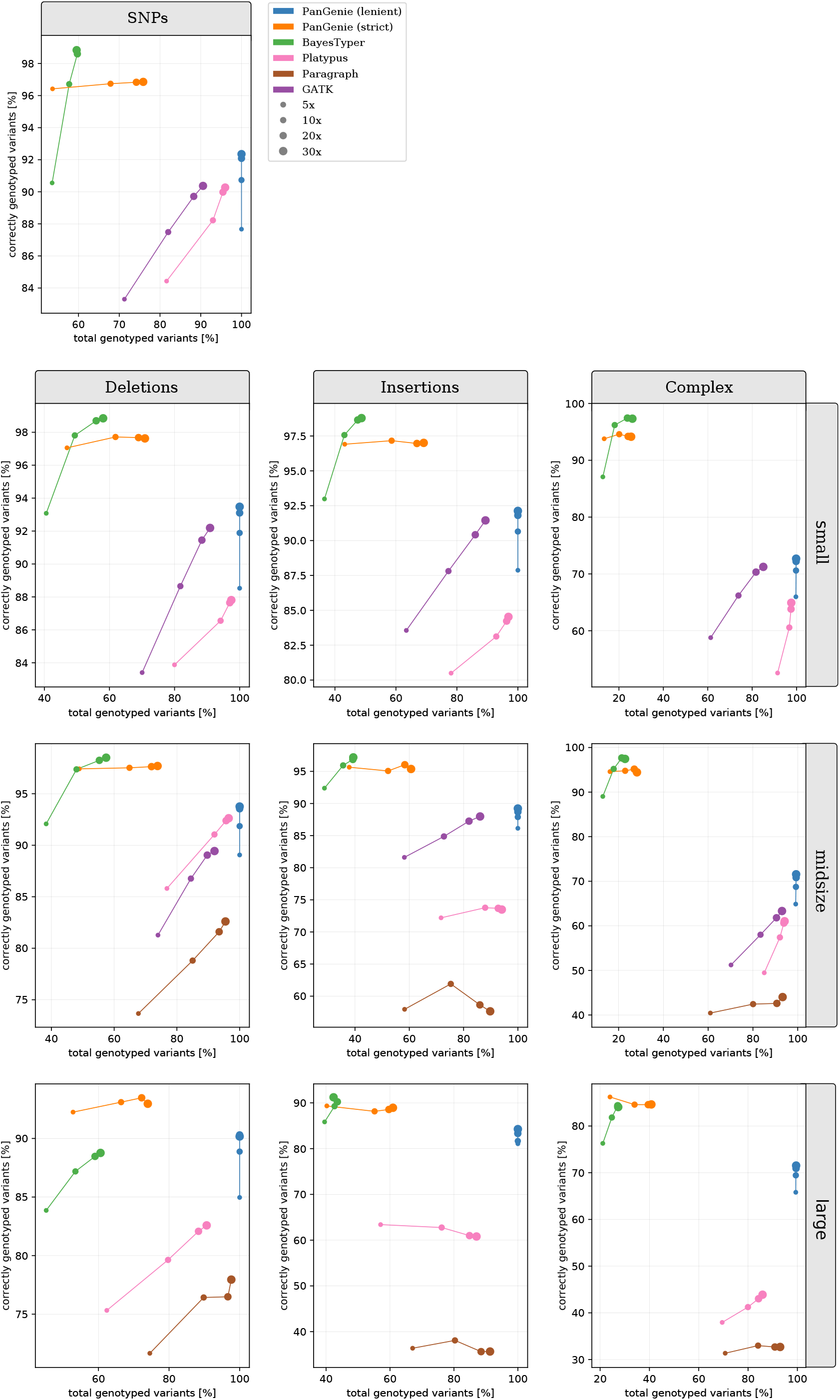
extended panel results for NA24385 in repeat regions. We ran PanGenie using the **extended** panel, BayesTyper, Paragraph, Platypus and GATK in order to re-genotype all callset variants located inside of repetitive STR/VNTR regions. Besides not applying any filter on the reported genotype qualities (“lenient”), we additionally report genotyping statistics for PanGenie when using “strict” filtering (genotype quality >= 200). Note that GATK was not run on large variants, and Paragraph was only run on midsize and large variants.

#### Results for the small panel

We provide the genotyping results that we obtained for all four samples using the small panel (8 haplotypes) in Figures 14-21. Experiments were run analogously to what we describe in Section 2.3.

**Figure 14:**
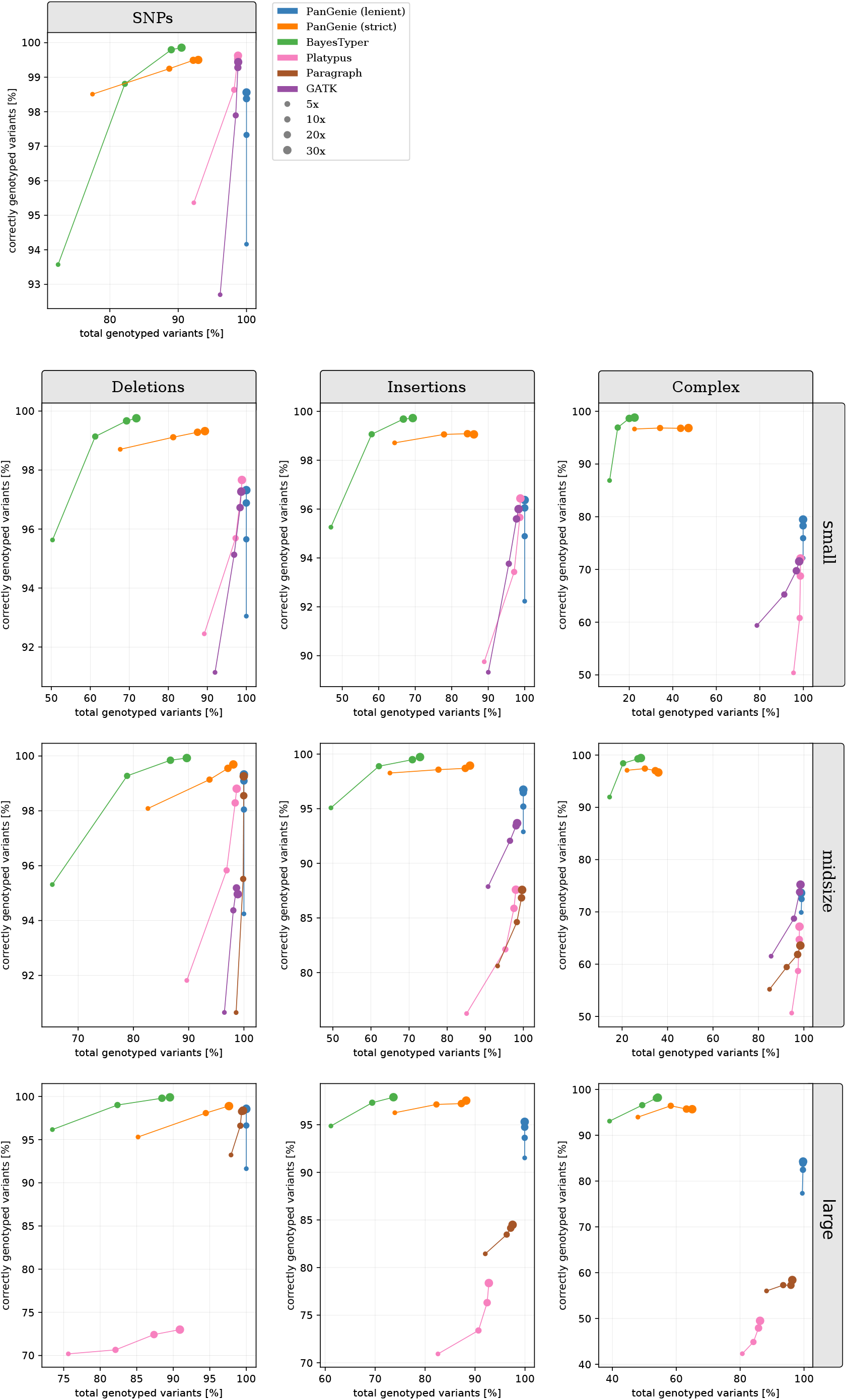
small panel results for HG00731 in non-repetitive regions. We ran PanGenie using the **small** panel, BayesTyper, Paragraph, Platypus and GATK in order to re-genotype all callset variants located outside of repetitive STR/VNTR regions. Besides not applying any filter on the reported genotype qualities (“lenient”), we additionally report genotyping statistics for PanGenie when using “strict” filtering (genotype quality >= 200). Note that GATK was not run on large variants, and Paragraph was only run on midsize and large variants.

**Figure 15:**
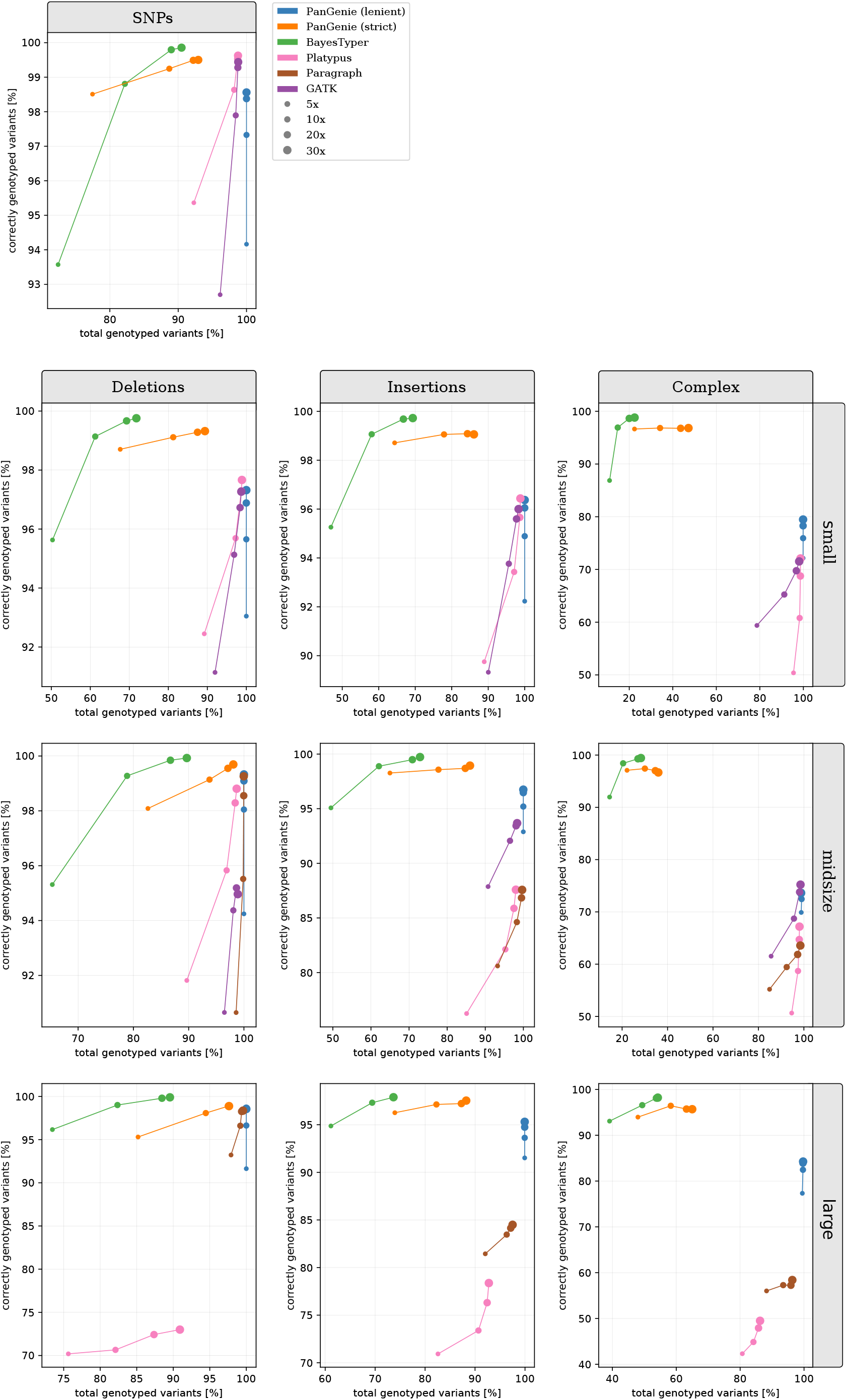
small panel results for HG00731 in repeat regions. We ran PanGenie using the **small** panel, BayesTyper, Paragraph, Platypus and GATK in order to re-genotype all callset variants located inside of repetitive STR/VNTR regions. Besides not applying any filter on the reported genotype qualities (“lenient”), we additionally report genotyping statistics for PanGenie when using “strict” filtering (genotype quality >= 200). Note that GATK was not run on large variants, and Paragraph was only run on midsize and large variants.

**Figure 16:**
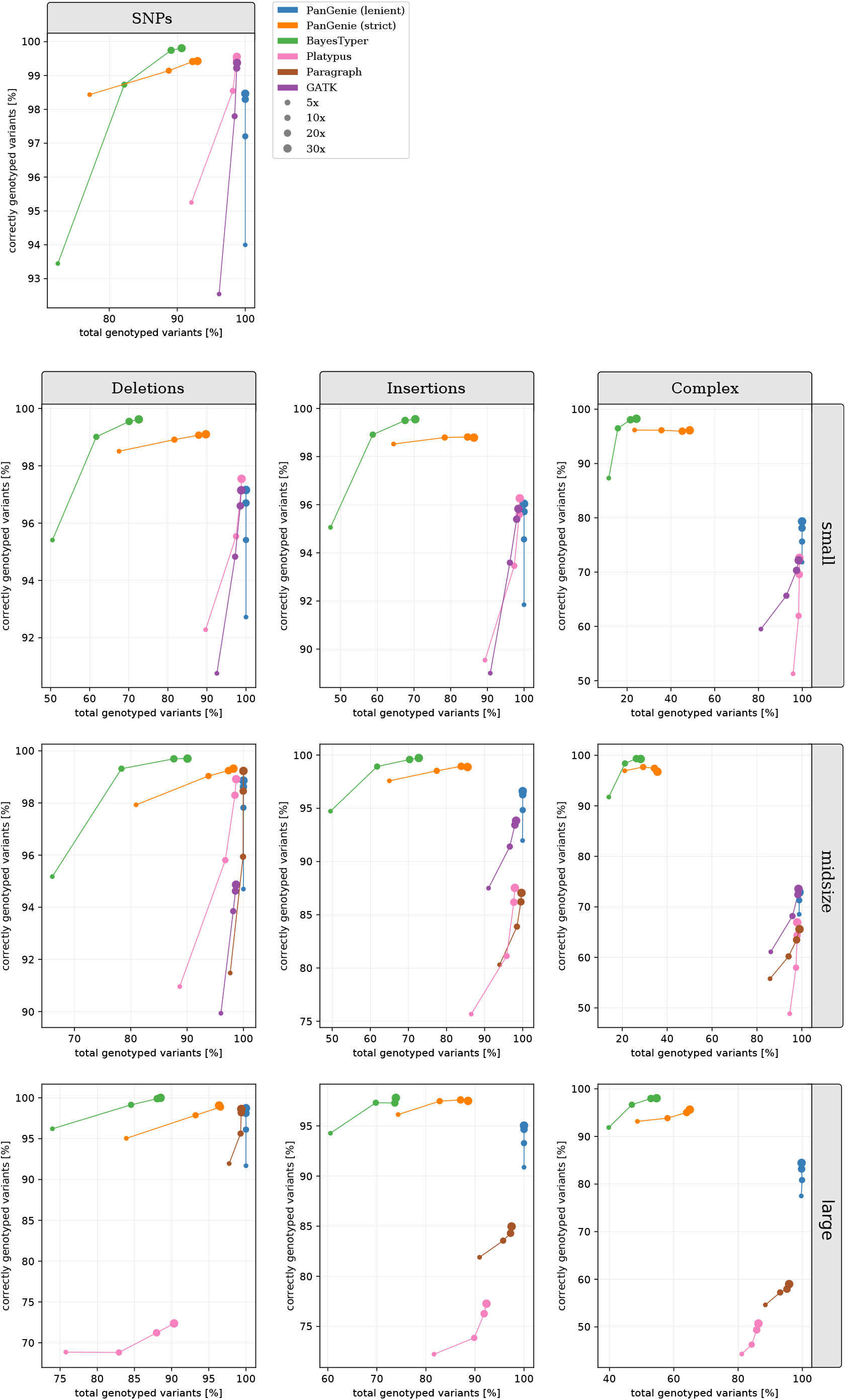
small panel results for HG00732 in non-repetitive regions. We ran PanGenie using the **small** panel, BayesTyper, Paragraph, Platypus and GATK in order to re-genotype all callset variants located outside of repetitive STR/VNTR regions. Besides not applying any filter on the reported genotype qualities (“lenient”), we additionally report genotyping statistics for PanGenie when using “strict” filtering (genotype quality >= 200). Note that GATK was not run on large variants, and Paragraph was only run on midsize and large variants.

**Figure 17:**
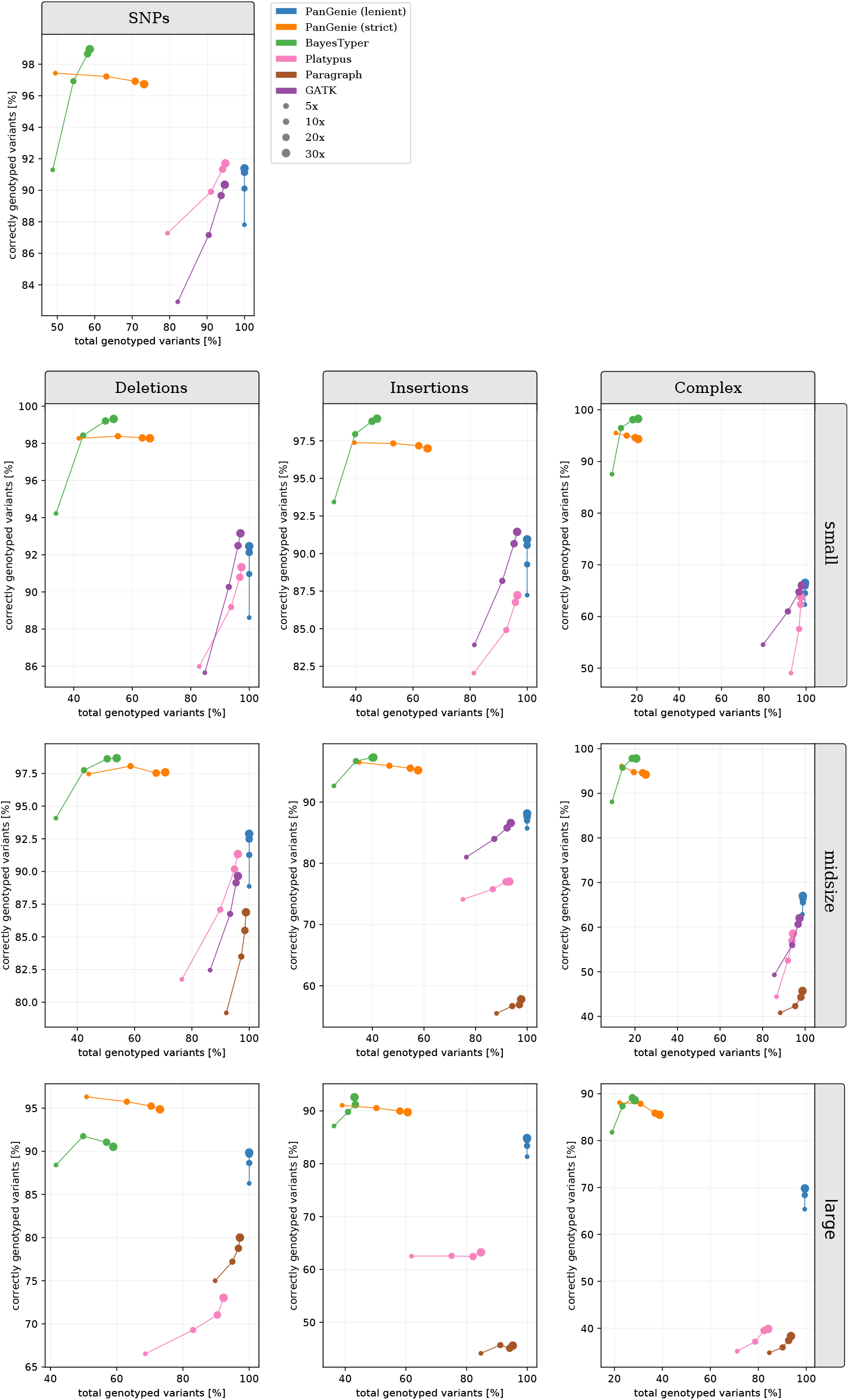
small panel results for HG00732 in repeat regions. We ran PanGenie using the **small** panel, BayesTyper, Paragraph, Platypus and GATK in order to re-genotype all callset variants located inside of repetitive STR/VNTR regions. Besides not applying any filter on the reported genotype qualities (“lenient”), we additionally report genotyping statistics for PanGenie when using “strict” filtering (genotype quality >= 200). Note that GATK was not run on large variants, and Paragraph was only run on midsize and large variants.

**Figure 18:**
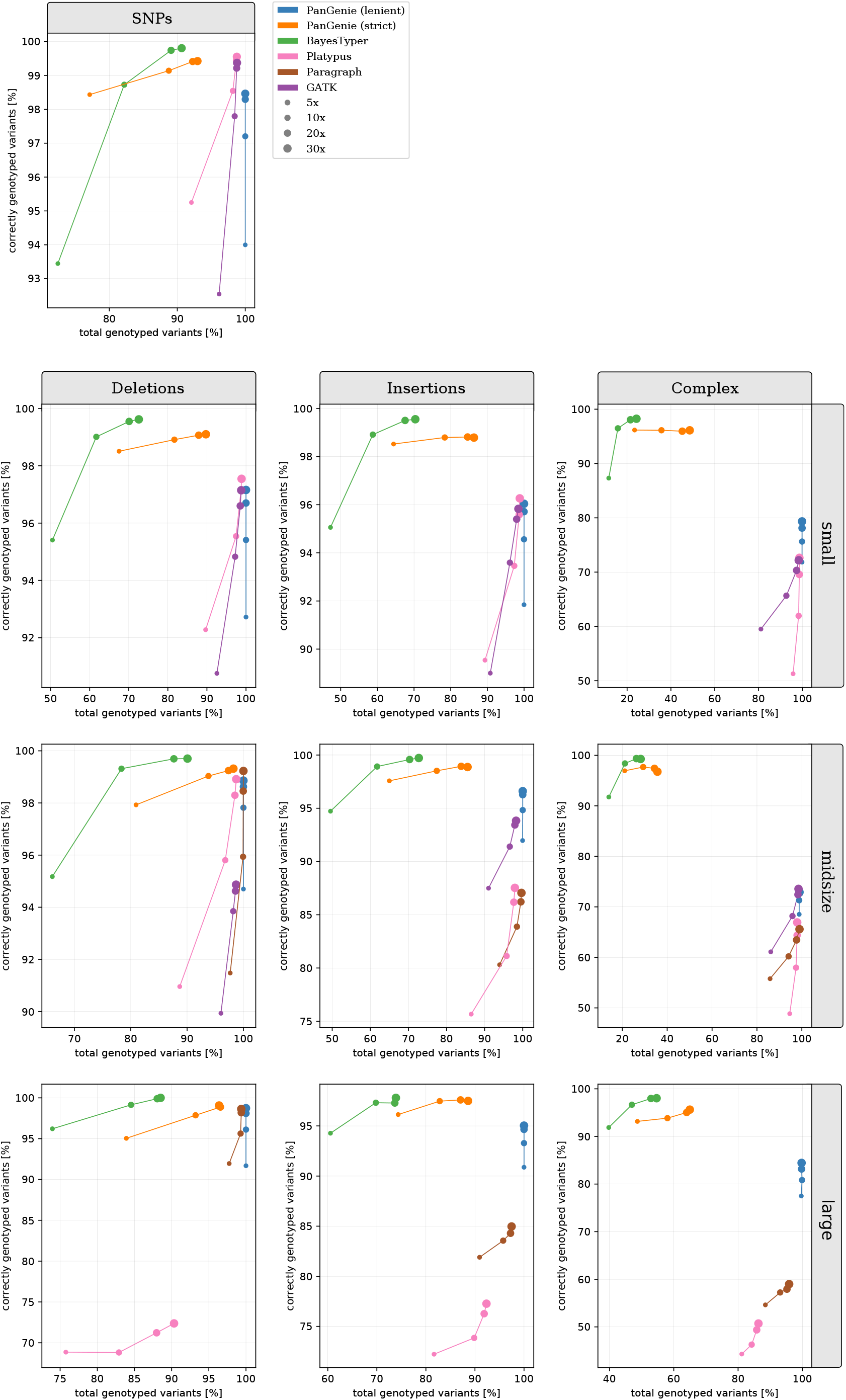
small panel results for NA12878 in non-repetitive regions. We ran PanGenie using the **small** panel, BayesTyper, Paragraph, Platypus and GATK in order to re-genotype all callset variants located outside of repetitive STR/VNTR regions. Besides not applying any filter on the reported genotype qualities (“lenient”), we additionally report genotyping statistics for PanGenie when using “strict” filtering (genotype quality >= 200). Note that GATK was not run on large variants, and Paragraph was only run on midsize and large variants.

**Figure 19:**
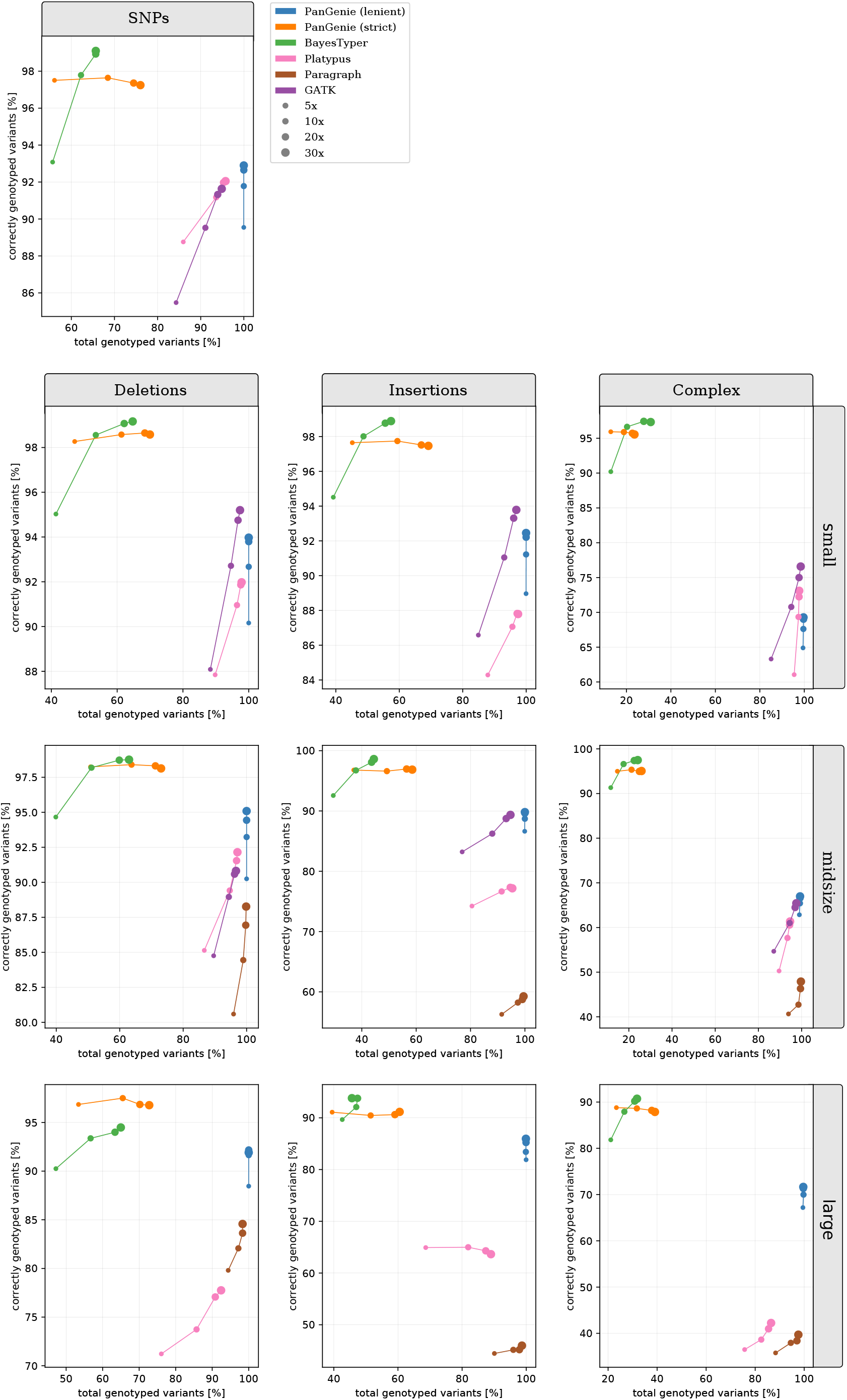
small panel results for NA12878 in repeat regions. We ran PanGenie using the **small** panel, BayesTyper, Paragraph, Platypus and GATK in order to re-genotype all callset variants located inside of repetitive STR/VNTR regions. Besides not applying any filter on the reported genotype qualities (“lenient”), we additionally report genotyping statistics for PanGenie when using “strict” filtering (genotype quality >= 200). Note that GATK was not run on large variants, and Paragraph was only run on midsize and large variants.

**Figure 20:**
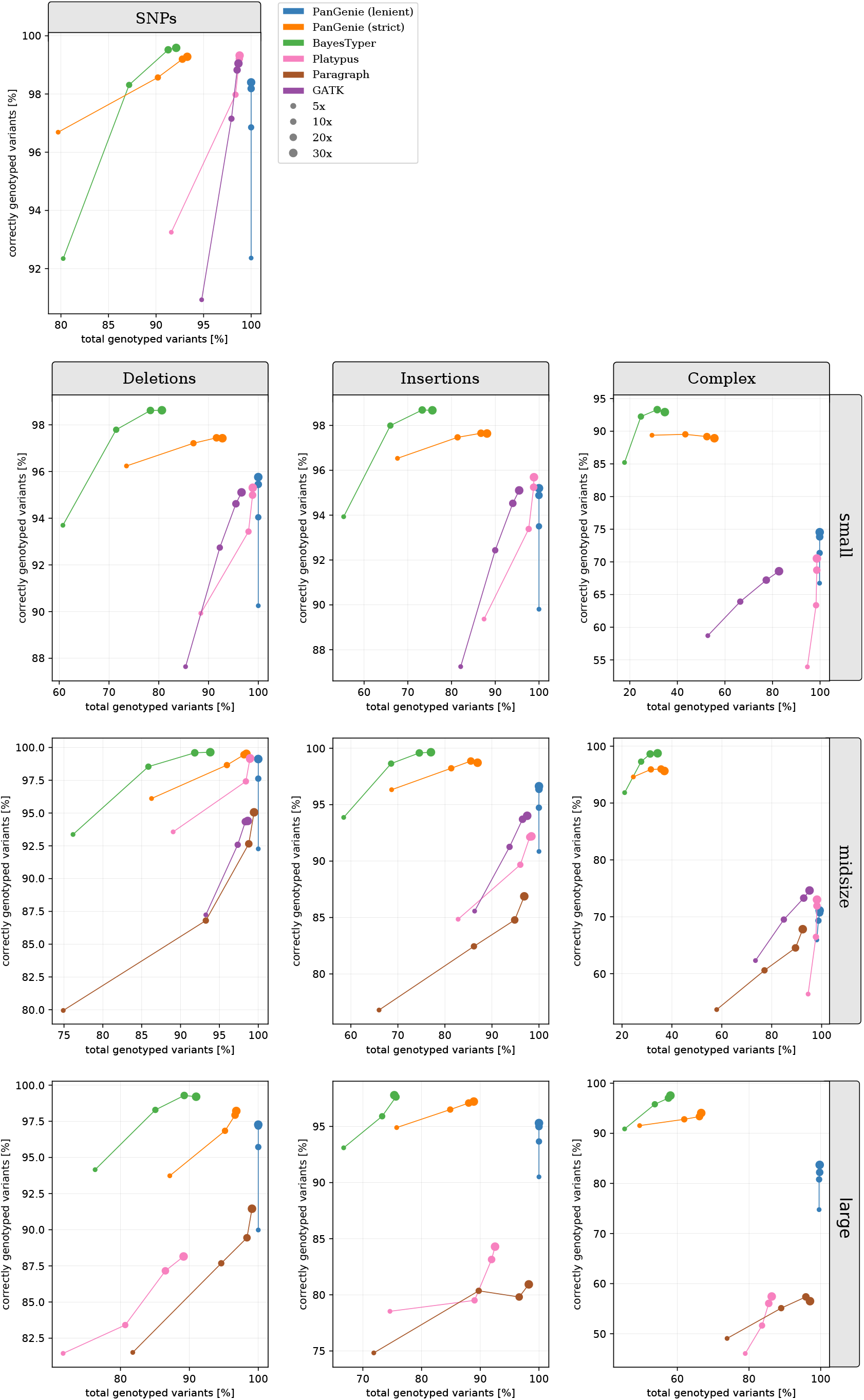
small panel results for NA24385 in non-repetitive regions. We ran PanGenie using the **small** panel, BayesTyper, Paragraph, Platypus and GATK in order to re-genotype all callset variants located outside of repetitive STR/VNTR regions. Besides not applying any filter on the reported genotype qualities (“lenient”), we additionally report genotyping statistics for PanGenie when using “strict” filtering (genotype quality>= 200). Note that GATK was not run on large variants, and Paragraph was only run on midsize and large variants.

**Figure 21:**
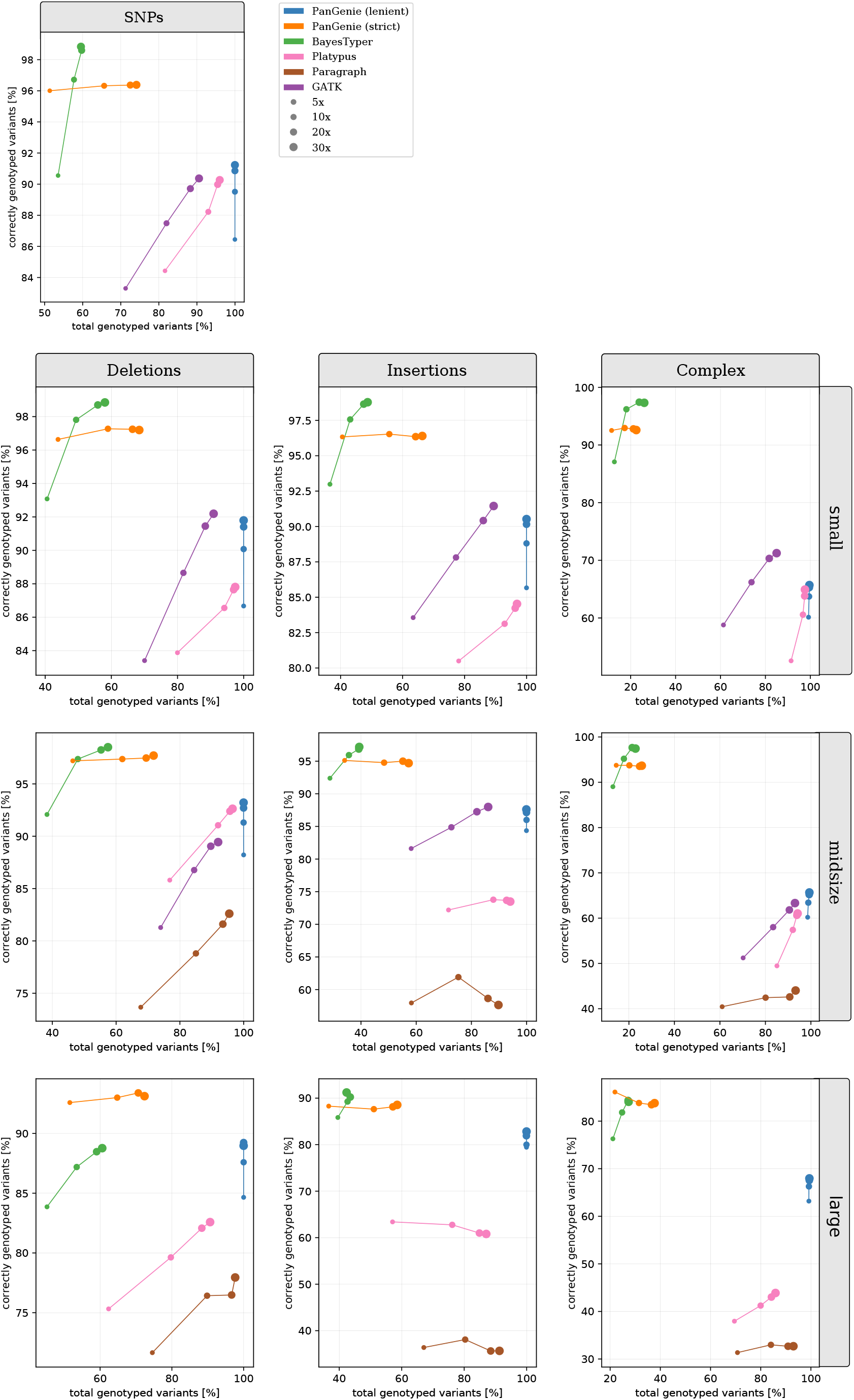
small panel results for NA24385 in repeat regions. We ran PanGenie using the **small** panel, BayesTyper, Paragraph, Platypus and GATK in order to re-genotype all callset variants located inside of repetitive STR/VNTR regions. Besides not applying any filter on the reported genotype qualities (“lenient”), we additionally report genotyping statistics for PanGenie when using “strict” filtering (genotype quality >= 200). Note that GATK was not run on large variants, and Paragraph was only run on midsize and large variants.

#### Runtimes

We measured the runtimes of all genotypers for the experiments described in Section 2.3 and show them in the table below. For all methods we measured the time needed to produce genotypes given the raw, unaligned sequencing reads. Therefore, runtimes for the mapping based approaches (Platypus, GATK, Paragraph) include the time that was needed to align the reads to the reference genome.

### Comparison to GIAB variants

We also evaluated our genotyping results for individual NA12878 taking the Genome in a Bottle (GIAB) small variant calls [43] as a ground truth. We determined all variants that the GIAB callset and our assembly-based VCF had in common and compared the genotype predictions made by the genotypers to the true genotypes. We only considered exact matches, that is, a variant was considered an overlap, if the positions and genotype alleles between both callsets where exactly identical. Figure 22 shows the results. Variants that were present in the GIAB callset but not our assembly calls, were treated as “untyped” when creating the plots.

**Figure 22:**
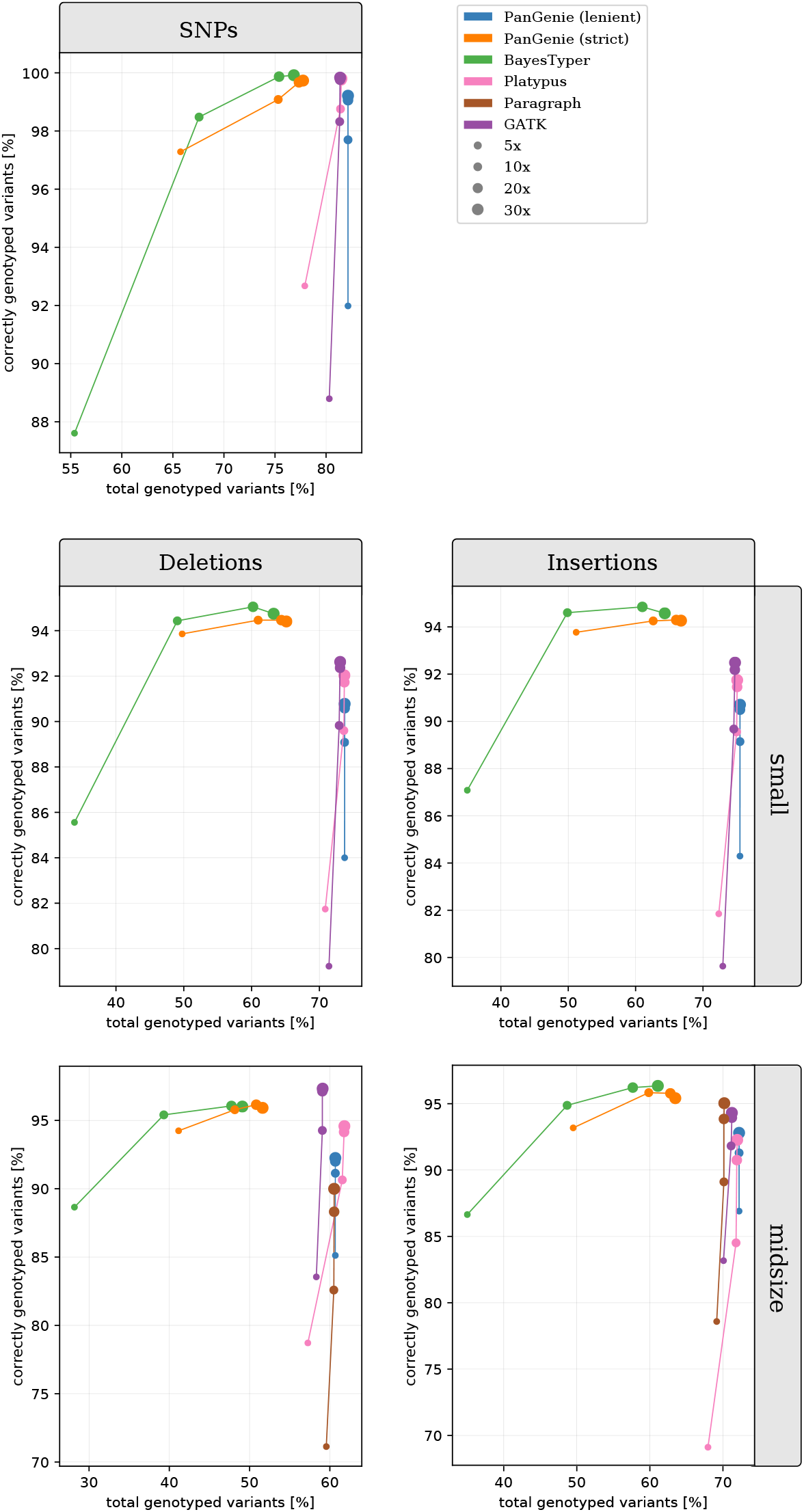
**Genotyping Performance** for sample **NA12878** on GIAB variants at different coverages. Evaluation of the genotypes produced by PanGenie, BayesTyper, Paragraph, Platypus and GATK for the variants overlapping with the Genome in a Bottle ground truth.

### Genotyping Larger Cohorts

Here we additionally show the results for the experiment described in Section 2.4 that we get when using strict filtering on the reported genotypes. At each biallelic variant position, only genotypes with a quality >= 200 were considered. Additionally, we skip if a genotype quality below that threshold was reported for more than ten samples. Results are shown in Figure 23.

**Figure 23:**
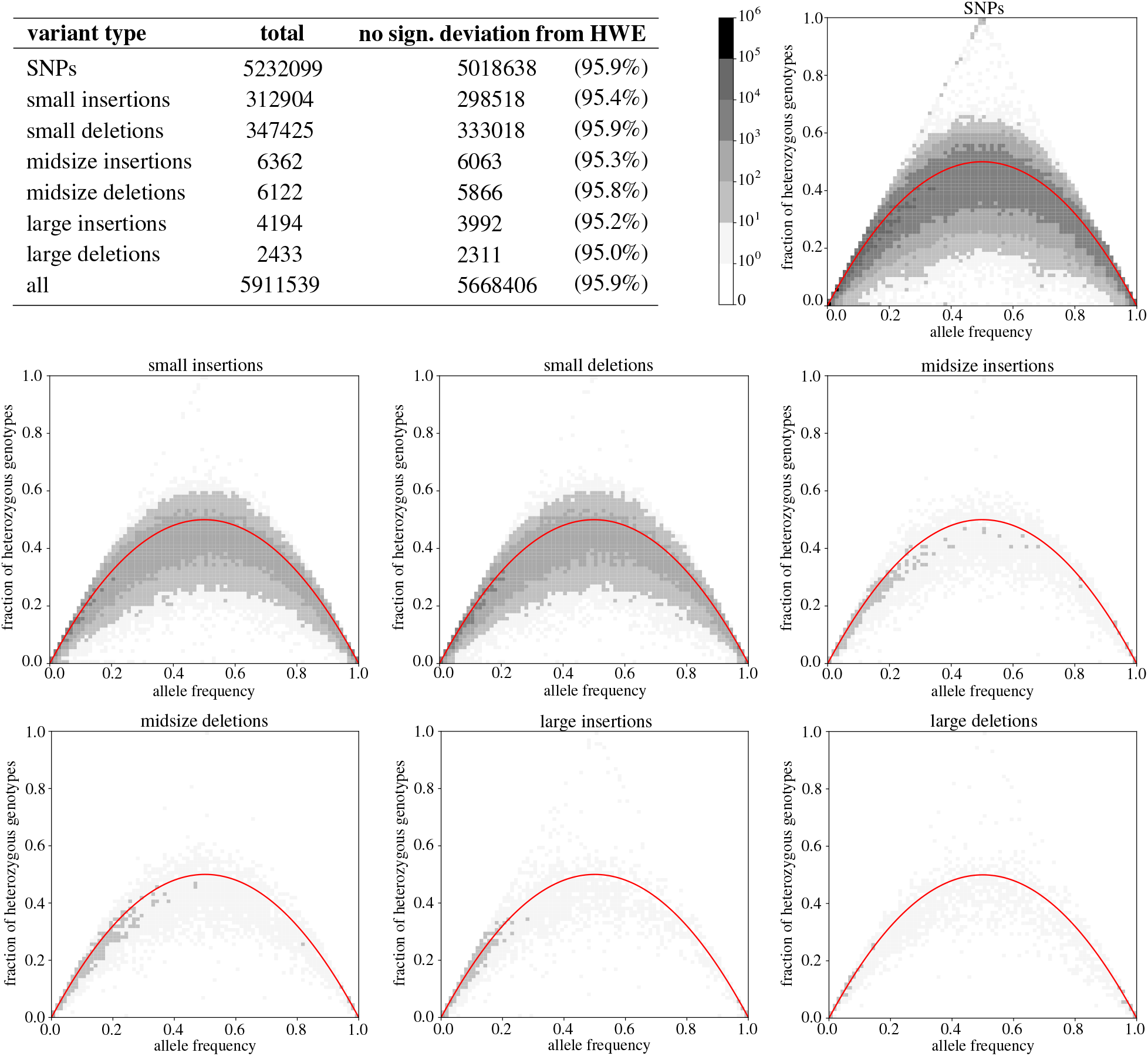
Genotyping larger cohorts (strict filtering). The table provides the amount of variants for which no significant deviation from Hardy-Weinberg Equilibrium was observed. Each plot shows the fraction of heterozygous genotypes at a variant position as a function of the allele frequency. The red curve shows what is expected according to Hardy-Weinberg equilibrium. Only genotypes with a quality of at least 200 were considered and positions with more than 10 low quality genotypes were skipped.

### Command lines used for genotyping

We used a VCF-file containing the variants detected from the haplotype-resolved assemblies as input variants for all genotyping tools and genotyped them based on short Illumina reads as described in Section 2.3. We ran BayesTyper (version v1.5) and PanGenie with default parameters using the raw, unaligned Illumina reads (FASTQ format) as input. For BayesTyper, we used the Snakemake pipeline provided in their repository (https://github.com/bioinformatics-centre/BayesTyper). PanGenie (https://bitbucket.org/jana_ebler/pangenie/src/master/, commit: `f46a9e5`) was run based on the command shown below,

~~~
PGGTyper -i reads.fq -v variants.vcf -r reference.fa -o pangenie-results -j 22 -t 22 -g
~~~

where `variants.vcf` refers to the input VCF file that contains the variants to be genotyped.

The remaining tools were provided with the aligned reads in BAM format, produced by mapping them to the reference genome using bwa. Platypus (version 0.8.1) was run in re-typing mode with additional options

~~~
–-source=variants.vcf, –-minPosterior=0 and –-getVariantsFromBAMs=0.
~~~

In order to run GATK (version 4.1.3.0), we first marked duplicates in our BAMs and then used HaplotypeCaller in re-typing mode in order to compute genotypes for the input variants using the command below. Note that we did not genotype large variants with GATK, therefore we removed them from the input VCF file prior to genotyping.

~~~
GATK HaplotypeCaller –reference reference.fa –-input reads.bam –-output GATK-results
~~~

~~~
–-minimum-mapping-quality 20 –-genotyping-mode GENOTYPE_GIVEN_ALLELES –-alleles
~~~

~~~
variants_no_large.vcf
~~~

In order to run Paragraph, we first computed the depth of the input BAM file using the command

~~~
/bin/idxdepth -b reads.bam -r reference.fasta -o depth.json
~~~

and prepared the Manifest file required for genotyping. In the next step, we used the command `bin/multigrmpy.py` with default parameters in order to genotype the input variants. Note that we removed all variants shorter than 20 bp from the input VCF before running Paragraph in order to only type midsize and large variants.

The complete pipeline used to run the evaluation including the commands used to run all tools can be found in this repository: https://bitbucket.org/jana_ebler/genotyping-experiments/src/master/

## Footnotes

1 https://www.genome.gov/news/news-release/NIH-funds-centers-for-advancing-sequence-of-human-genome-reference

